# Alveolitis in severe SARS-CoV-2 pneumonia is driven by self-sustaining circuits between infected alveolar macrophages and T cells

**DOI:** 10.1101/2020.08.05.238188

**Authors:** Rogan A. Grant, Luisa Morales-Nebreda, Nikolay S. Markov, Suchitra Swaminathan, Estefany R. Guzman, Darryl A. Abbott, Helen K. Donnelly, Alvaro Donayre, Isaac A. Goldberg, Zasu M. Klug, Nicole Borkowski, Ziyan Lu, Hermon Kihshen, Yuliya Politanska, Lango Sichizya, Mengjia Kang, Ali Shilatifard, Chao Qi, A. Christine Argento, Jacqueline M. Kruser, Elizabeth S. Malsin, Chiagozie O. Pickens, Sean Smith, James M. Walter, Anna E. Pawlowski, Daniel Schneider, Prasanth Nannapaneni, Hiam Abdala-Valencia, Ankit Bharat, Cara J. Gottardi, GR Scott Budinger, Alexander V. Misharin, Benjamin D. Singer, Richard G. Wunderink, for The NU SCRIPT Study Investigators

**Author notes:** These authors contributed equally to this work and will list themselves first on their CVs (Rogan A. Grant, Luisa Morales-Nebreda, Nikolay S. Markov). Corresponding authors contributed equally to this work and listed in alphabetical order.

## Abstract

Some patients infected with Severe Acute Respiratory Syndrome Coronavirus-2 (SARS-CoV-2) develop severe pneumonia and the acute respiratory distress syndrome (ARDS) [1]. Distinct clinical features in these patients have led to speculation that the immune response to virus in the SARS-CoV-2-infected alveolus differs from other types of pneumonia [2]. We collected bronchoalveolar lavage fluid samples from 86 patients with SARS-CoV-2-induced respiratory failure and 252 patients with known or suspected pneumonia from other pathogens and subjected them to flow cytometry and bulk transcriptomic profiling. We performed single cell RNA-Seq in 5 bronchoalveolar lavage fluid samples collected from patients with severe COVID-19 within 48 hours of intubation. In the majority of patients with SARS-CoV-2 infection at the onset of mechanical ventilation, the alveolar space is persistently enriched in alveolar macrophages and T cells without neutrophilia. Bulk and single cell transcriptomic profiling suggest SARS-CoV-2 infects alveolar macrophages that respond by recruiting T cells. These T cells release interferon-gamma to induce inflammatory cytokine release from alveolar macrophages and further promote T cell recruitment. Our results suggest SARS-CoV-2 causes a slowly unfolding, spatially-limited alveolitis in which alveolar macrophages harboring SARS-CoV-2 transcripts and T cells form a positive feedback loop that drives progressive alveolar inflammation.

This manuscript is accompanied by an online resource: https://www.nupulmonary.org/covid-19/

**One sentence summary:** SARS-CoV-2-infected alveolar macrophages form positive feedback loops with T cells in patients with severe COVID-19.

## Introduction

Most patients infected by Severe Acute Respiratory Syndrome Coronavirus-2 (SARS-CoV-2) suffer no or only mild symptoms. A small minority of patients develop severe Coronavirus Disease 2019 (COVID-19), and these patients account for almost all of the morbidity and mortality associated with the infection [1]. Reported rates of mortality in patients with severe disease range between 20–40% [3–5]. The relatively high mortality among patients with severe SARS-CoV-2 pneumonia reported in some centers, combined with a systemic inflammatory response that is severe in some patients, have led to speculation that the pathobiology of SARS-CoV-2 pneumonia is distinct from other respiratory viral and bacterial pathogens [2].

We obtained bronchoalveolar lavage (BAL) fluid from 86 patients with respiratory failure secondary to severe SARS-CoV-2 pneumonia and compared them with BAL specimens from 252 patients with pneumonia secondary to other pathogens and intubated patients without pneumonia that were prospectively collected before and during the pandemic. For many patients we were able to obtain samples within 48 hours of intubation and sequentially over the course of the illness, allowing us to gain insights about the early pathogenesis of COVID-19-induced acute respiratory distress syndrome (ARDS). We profiled BAL samples using multicolor flow cytometry to identify CD4+ and CD8+ T cells, monocytes, mature and immature alveolar macrophages, and neutrophils. We also performed bulk transcriptomic profiling of flow cytometry-sorted alveolar macrophages in a subset of patients with confirmed COVID-19. Additionally, we performed single cell RNA-Seq on BAL fluid collected less than 48 hours after intubation from 5 patients with severe SARS-CoV-2 pneumonia.

Our results suggest that, unlike other respiratory pathogens such as influenza viruses and bacteria, the alveolar space of SARS-CoV-2-infected patients is populated by alveolar macrophages and T cells without neutrophilia. We also detected positive and negative SARS-CoV-2 viral transcripts in alveolar macrophages in the absence of *ACE2* expression, suggestive of active viral replication in alveolar macrophages. Alveolar macrophages harboring SARS-CoV-2 transcripts are activated and express *IL1B*, a master regulator of inflammation, and several chemokines known to drive recruitment of T cells. These same alveolar macrophages show an enhanced response to interferon-gamma (IFNγ) generated by T cells. Our results suggest that SARS-CoV-2 infection results in a slowly progressing and spatially limited alveolitis, in which alveolar macrophages harboring SARS-CoV-2 and IFNγ-producing T cells form self-sustaining circuits that drive alveolar inflammation. We further suggest that alveolar macrophages may be a nonspecific reservoir for the SARS-CoV-2 virus, acting as a Trojan horse to enable immune evasion and viral dissemination throughout the lung.

## Results

### Demographics

Samples were collected from patients enrolled in the Successful Clinical Response in Pneumonia Therapy (SCRIPT) systems biology center, a NIAID-funded observational study of patients with severe pneumonia, defined as those requiring mechanical ventilation. From March 13 to June 15, 2020, we prospectively enrolled 86 of the 256 patients with SARS-CoV-2-induced pneumonia and respiratory failure requiring mechanical ventilation in our ICU at Northwestern Medicine, Chicago (**Figure S1a, b; Table 1**). We obtained sequential samples in the majority of patients (**Figure 1a**). We compared these patients with 252 mechanically ventilated patients with known or clinically-suspected pneumonia enrolled in the two years before and during the pandemic. Five critical care physicians (RGW, JMK, BDS, COP, JMW) retrospectively adjudicated patients as COVID-19 pneumonia, non-COVID-19 viral pneumonia, pneumonia secondary to other pathogens or non-pneumonia controls (intubated for reasons other than pneumonia) according to a standardized adjudication procedure (see **Methods**). Patients were designated as viral pneumonia if a multiplex PCR for respiratory viruses obtained by nasopharyngeal swab or BAL was positive in the absence of an alternative diagnosis based on quantitative culture of BAL fluid or multiplex PCR. A complete list of viral and bacterial pathogens identified as the cause of pneumonia is in **Table 2**. For some patients without COVID-19 the diagnosis of pneumonia was made based on clinical suspicion, radiographic findings and response to antimicrobial therapy in the absence of an identified pathogen. For all patients with COVID-19, samples were collected from regions of greatest chest radiograph abnormality by a critical care physician using a disposable bronchoscope. The majority of samples prior to the pandemic were collected by respiratory therapists using non-bronchoscopic bronchoalveolar lavage (NBBAL) with the catheter directed to the most radiographically affected lung [6].

**Table 1:** Demographics of the SCRIPT cohort, grouped by diagnosis.

|  | Diagnosis: COVID-19 (N = 74) | Diagnosis: Non-Pneumonia Control (N = 39) | Diagnosis: Other Pneumonia (N = 147) | Diagnosis: Other Viral Pneumonia (N = 37) |
| --- | --- | --- | --- | --- |
| **Ethnicity** |  |  |  |  |
| Hispanic or Latino | 33/74 (44.6%) | 6/39 (15.4%) | 10/147 (6.8%) | 8/37 (21.6%) |
| Not Hispanic or Latino | 33/74 (44.6%) | 30/39 (76.9%) | 127/147 (86.4%) | 26/37 (70.3%) |
| Unknown or Not Reported | 8/74 (10.8%) | 3/39 (7.7%) | 10/147 (6.8%) | 3/37 (8.1%) |
| **Race** |  |  |  |  |
| American Indian/Alaska Native | 0/74 (0.0%) | 0/39 (0.0%) | 0/147 (0.0%) | 1/37 (2.7%) |
| Asian | 1/74 (1.4%) | 2/39 (5.1%) | 6/147 (4.1%) | 1/37 (2.7%) |
| Black/African American | 16/74 (21.6%) | 6/39 (15.4%) | 32/147 (21.8%) | 7/37 (18.9%) |
| Unknown or Not Reported | 25/74 (33.8%) | 5/39 (12.8%) | 11/147 (7.5%) | 5/37 (13.5%) |
| White | 32/74 (43.2%) | 26/39 (66.7%) | 98/147 (66.7%) | 23/37 (62.2%) |
| **Age** |  |  |  |  |
| Minimum | 21 | 19 | 22 | 34 |
| Median (IQR) | 61.00 (47.25, 69.00) | 64.00 (45.00, 72.00) | 66.00 (51.50, 73.00) | 58.00 (52.00, 69.00) |
| Mean (sd) | 57.97 ± 14.65 | 59.31 ± 18.53 | 62.15 ± 16.67 | 60.03 ± 13.81 |
| Maximum | 86 | 90 | 120 | 88 |
| **Sex** |  |  |  |  |
| Female | 24/74 (32.4%) | 17/39 (43.6%) | 60/147 (40.8%) | 16/37 (43.2%) |
| Male | 50/74 (67.6%) | 22/39 (56.4%) | 87/147 (59.2%) | 21/37 (56.8%) |
| **Discharge Status** |  |  |  |  |
| Deceased | 16/74 (21.6%) | 15/39 (38.5%) | 51/147 (34.7%) | 12/37 (32.4%) |
| Discharged | 34/74 (45.9%) | 17/39 (43.6%) | 28/147 (19.0%) | 9/37 (24.3%) |
| Inpatient Facility | 22/74 (29.7%) | 7/39 (17.9%) | 67/147 (45.6%) | 16/37 (43.2%) |
| Other | 2/74 (2.7%) | 0/39 (0.0%) | 1/147 (0.7%) | 0/37 (0.0%) |
| **Smoking Status** |  |  |  |  |
| Current Smoker | 0/74 (0.0%) | 6/39 (15.4%) | 21/147 (14.3%) | 5/37 (13.5%) |
| Never Smoker | 38/74 (51.4%) | 23/39 (59.0%) | 66/147 (44.9%) | 18/37 (48.6%) |
| Past Smoker | 12/74 (16.2%) | 9/39 (23.1%) | 58/147 (39.5%) | 14/37 (37.8%) |
| Unknown or Not Reported | 24/74 (32.4%) | 1/39 (2.6%) | 2/147 (1.4%) | 0/37 (0.0%) |
| **Diagnosis** |  |  |  |  |
| COVID-19 | 74/74 (100.0%) | 0/39 (0.0%) | 0/147 (0.0%) | 0/37 (0.0%) |
| Non-Pneumonia Control | 0/74 (0.0%) | 39/39 (100.0%) | 0/147 (0.0%) | 0/37 (0.0%) |
| Other Pneumonia | 0/74 (0.0%) | 0/39 (0.0%) | 147/147 (100.0%) | 0/37 (0.0%) |
| Other Viral Pneumonia | 0/74 (0.0%) | 0/39 (0.0%) | 0/147 (0.0%) | 37/37 (100.0%) |
| **Days of Intubation** |  |  |  |  |
| Minimum | 0 | 0 | 0 | 0 |
| Median (IQR) | 11.50 (1.25, 25.00) | 2.00 (1.00, 5.00) | 4.00 (1.00, 12.25) | 5.00 (2.00, 8.00) |
| Mean (sd) | 15.72 ± 15.08 | 3.92 ± 5.01 | 9.91 ± 14.60 | 9.00 ± 12.67 |
| Maximum | 61 | 23 | 110 | 63 |
| Unknown | 0/74 (0.0%) | 2/39 (5.1%) | 11/147 (7.5%) | 4/37 (10.8%) |
| **Length of ICU Stay** |  |  |  |  |
| Minimum | 3 | 1 | 1 | 1 |
| Median (IQR) | 17.00 (10.00, 29.75) | 5.00 (3.00, 9.50) | 9.00 (5.00, 18.00) | 11.00 (4.00, 18.00) |
| Mean (sd) | 21.24 ± 13.64 | 8.08 ± 7.74 | 12.88 ± 12.40 | 12.05 ± 8.64 |
| Maximum | 63 | 38 | 84 | 33 |
| Unknown | 0/74 (0.0%) | 0/39 (0.0%) | 2/147 (1.4%) | 0/37 (0.0%) |
| **BMI** |  |  |  |  |
| Minimum | 22.30 | 16.56 | 12.46 | 17.34 |
| Median (IQR) | 31.25 (27.64, 37.22) | 27.78 (24.65, 33.73) | 25.52 (21.67, 31.64) | 26.81 (22.70, 31.87) |
| Mean (sd) | 33.58 ± 9.00 | 30.53 ± 9.45 | 27.30 ± 8.54 | 28.99 ± 8.99 |
| Maximum | 81.81 | 70.60 | 70.25 | 60.26 |
| Unknown | 0/74 (0.0%) | 1/39 (2.6%) | 0/147 (0.0%) | 0/37 (0.0%) |
| **SOFA** |  |  |  |  |
| Minimum | 1.83 | 1.75 | 1.00 | 2.00 |
| Median (IQR) | 6.86 (5.42, 8.16) | 7.00 (4.73, 10.47) | 6.40 (4.40, 8.89) | 6.14 (3.86, 8.17) |
| Mean (sd) | 7.23 ± 2.51 | 7.45 ± 3.14 | 6.98 ± 3.46 | 6.26 ± 2.84 |
| Maximum | 14.84 | 13.75 | 18.50 | 13.17 |
| Unknown | 0/74 (0.0%) | 0/39 (0.0%) | 2/147 (1.4%) | 0/37 (0.0%) |
| **APS** |  |  |  |  |
| Minimum | 33.00 | 28.25 | 21.20 | 26.33 |
| Median (IQR) | 70.27 (57.44, 80.07) | 62.13 (46.03, 82.14) | 64.63 (50.72, 80.67) | 70.60 (51.20, 78.77) |
| Mean (sd) | 69.18 ± 17.32 | 65.83 ± 22.19 | 66.42 ± 22.71 | 66.98 ± 19.73 |
| Maximum | 103.36 | 115.50 | 161.00 | 106.50 |
| Unknown | 3/74 (4.1%) | 0/39 (0.0%) | 2/147 (1.4%) | 0/37 (0.0%) |
| **Outcome** |  |  |  |  |
| Deceased | 16/74 (21.6%) | 15/39 (38.5%) | 51/147 (34.7%) | 12/37 (32.4%) |
| Discharged | 58/74 (78.4%) | 24/39 (61.5%) | 96/147 (65.3%) | 25/37 (67.6%) |

**Table 2:** Pneumonia-causing pathogens detected in the SCRIPT cohort.

|  |  |
| --- | --- |
| 1 *Achromobacter denitrificans* | 25 *Haemophilus species, not influenzae* |
| 2 *Achromobacter species* | 26 *Hafnia alvei* |
| 3 *Achromobacter xylosoxidans* | 27 *Klebsiella oxytoca* |
| 4 *Acinetobacter baumannii* | 28 *Klebsiella pneumoniae* |
| 5 *Acinetobacter ursingii* | 29 *Staphylococcus aureus* |
| 6 *Beta Hemolytic Streptococci, Group C* | 30 *Moraxella catarrhalis* |
| 7 *Beta Hemolytic Streptococci, Group F* | 31 *Morganella morganii* |
| 8 *Beta Hemolytic Streptococci, Group G* | 32 *Neisseria meningitidis* |
| 9 *Beta Hemolytic Streptococci, not Group A, B, C, D, F, or G* | 33 *Neisseria species* |
| 10 *Burkholderia cepacia complex* | 34 *Pantoea species* |
| 11 *Capnocytophaga species* | 35 *Proteus mirabilis* |
| 12 *Chryseobacterium indologenes* | 36 *Providencia stuartii* |
| 13 *Citrobacter freundii* | 37 *Pseudomonas aeruginosa* |
| 14 *Citrobacter koseri* | 38 *Pseudomonas stutzeri* |
| 15 *Citrobacter youngae* | 39 *Raoultella ornithinolytica* |
| 16 *Elizabethkingia meningoseptica* | 40 *Salmonella species* |
| 17 *Enterobacter aerogenes* | 41 *Serratia marcescens* |
| 18 *Enterobacter cloacae complex* | 42 *Staphylococcus coagulase negative* |
| 19 *Enterococcus faecalis* | 43 *Stenotrophomonas maltophilia* |
| 20 *Enterococcus faecium* | 44 *Streptococcus agalactiae (Group B)* |
| 21 *Enterococcus species* | 45 *Streptococcus pneumoniae* |
| 22 *Escherichia coli* | 46 *Streptococcus pseudopneumoniae* |
| 23 *Granulicatella species* | 47 *Streptococcus pyogenes (Group A)* |
| 24 *Haemophilus influenzae* | 48 *Viridans streptococcus* |

**Figure 1.**
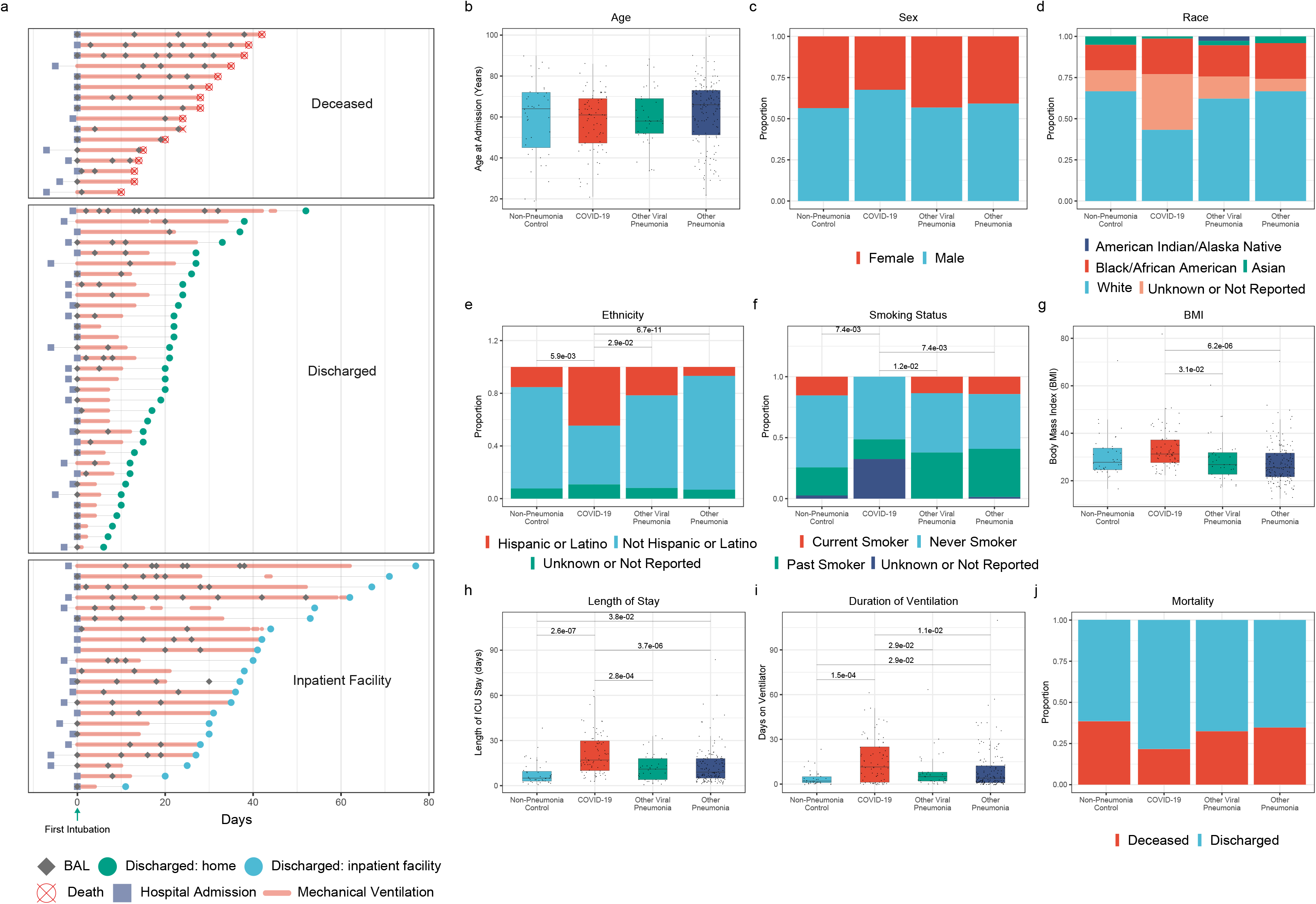
Demographics of the SCRIPT cohort. We compared BAL fluid obtained sequentially from 86 patients with severe SARS-CoV-2 pneumonia requiring mechanical ventilation with 39 patients with confirmed pneumonia secondary to other respiratory viruses (other viral pneumonia) and 159 patients with non-viral pneumonia (other pneumonia) and 41 mechanically ventilated patients without pneumonia undergoing BAL (non-pneumonia controls). **a**. Timing of hospital admission (square), BAL collection (diamonds) and length of mechanical ventilation (bold red line) and hospital stay (thin grey line) in patients with severe COVID-19 grouped by outcomes (crossed open red circles – death; green circles – discharged home; light blue circles – discharged to inpatient facility). Day 0 is defined as the day of the first intubation. An average of 1.63 (range 1–9) BAL samples were collected per patient. **b**. Distribution of patient ages. Differences not significant by pairwise t-test with FDR correction. **c**. Sex. Proportions of females (red) and males (blue) were similar between groups (pairwise Chi-Square tests of proportions with continuity and FDR correction). **d**. Self-reported race. Proportions were similar between groups (pairwise Chi-Square tests of proportions with continuity and FDR correction). **e**. Self-reported ethnicity. A significant increase in the proportion of individuals identifying as Hispanic or Latino was observed in the COVID-19 cohort, relative to all controls (q < 0.05, pairwise Chi-Square tests of proportions with continuity and FDR correction). **f**. Self-reported smoking status. Significantly fewer active smokers were observed in the COVID-19 cohort, as compared with all control groups (q < 0.05, pairwise Chi-Square tests of proportions with continuity and FDR correction). **g**. Body mass index (BMI). BMI was elevated in the COVID-19 cohort relative to patients with viral pneumonia, and other pneumonia, but not to non-pneumonia controls (q < 0.05, t-test with FDR correction). **h**. ICU length of stay. ICU length of stay was longer in the COVID-19 cohort, relative to all control cohorts (pairwise t-tests with FDR correction). **i**. Duration of mechanical ventilation. The duration of mechanical ventilation was significantly increased in the COVID-19 cohort relative only to non-pneumonia controls (pairwise t-tests with FDR correction). **j**. Mortality in patients with COVID-19 was similar to patients with other respiratory viruses, non-viral pneumonia and non-pneumonia controls. Differences are not significant (q > 0.05) unless otherwise noted.

Compared with patients with known or suspected pneumonia requiring mechanical ventilation, patients with severe SARS-CoV-2 pneumonia were similar in age, race, and sex, but had a significantly higher body mass index and a lower proportion of patients were current smokers, compared with historic control populations diagnosed with other viral pneumonias, non-viral pneumonia, or non-pneumonia control patients (**Figure 1b-g**). Patients with severe SARS-CoV-2 pneumonia required longer periods of ventilation and had longer lengths of stay in the ICU compared with all pneumonia and non-pneumonia controls (**Figure 1h-i).** Severity of illness estimated using the sequential organ failure assessment (SOFA) score and the acute physiology score (APS) was similar in patients with SARS-CoV-2 pneumonia compared with other pneumonia and was comparable to that observed in a recent study of ARDS [7] (**Supplemental Figure S1c** and **Table 1**). Nevertheless, mortality was significantly lower (q = 0.046, Chi-Square tests of proportions) in patients with SARS-CoV-2 pneumonia, compared with the entire cohort (22% vs 35%), but was not significantly different as compared with any individual comparison group (**Figure 1j**).

### The BAL fluid from patients with SARS-CoV-2 pneumonia is dominated by T cells

To define the immune cell profile over the course of severe SARS-CoV-2-induced pneumonia, we analyzed 116 samples from 61 patients with confirmed COVID-19 in our cohort. We compared those samples with 228 samples from 179 patients with severe pneumonia secondary to other causes. We used multicolor flow cytometry with a panel of markers sufficient to identify neutrophils, CD4+ and CD8+ T cell subsets, macrophages and monocytes in the BAL fluid (**Figure S2a,b**) [6,8]. We first focused analysis on patients who had a sample collected within 48 hours of intubation, comparing 29 samples from 28 patients with confirmed COVID-19 to 89 samples from 85 patients with severe pneumonia secondary to other causes. We found that, despite a diagnosis of severe ARDS requiring mechanical ventilation, only 34.4% of patients with severe COVID-19 exhibited neutrophilia in BAL fluid within 48 hours of intubation, defined as a neutrophil frequency over 50% (**Figure 2a–c**). Instead, we found that in patients with severe SARS-CoV-2 pneumonia, the alveolar space was significantly enriched for both CD4+ and CD8+ T cells and monocytes (**Figure 2a,b**). This observation significantly differs from patients with pneumonia secondary to other respiratory viruses, bacteria or mechanically ventilated non-pneumonia controls [8]. This pattern was not due to the differences in co-infections between different types of pneumonia (**Figure 2d**). Collectively, these findings suggest that unusual factors related to SARS-CoV-2 pneumonia result in the recruitment of T cells and monocytes, rather than neutrophils, to the infected alveolus.

**Figure 2.**
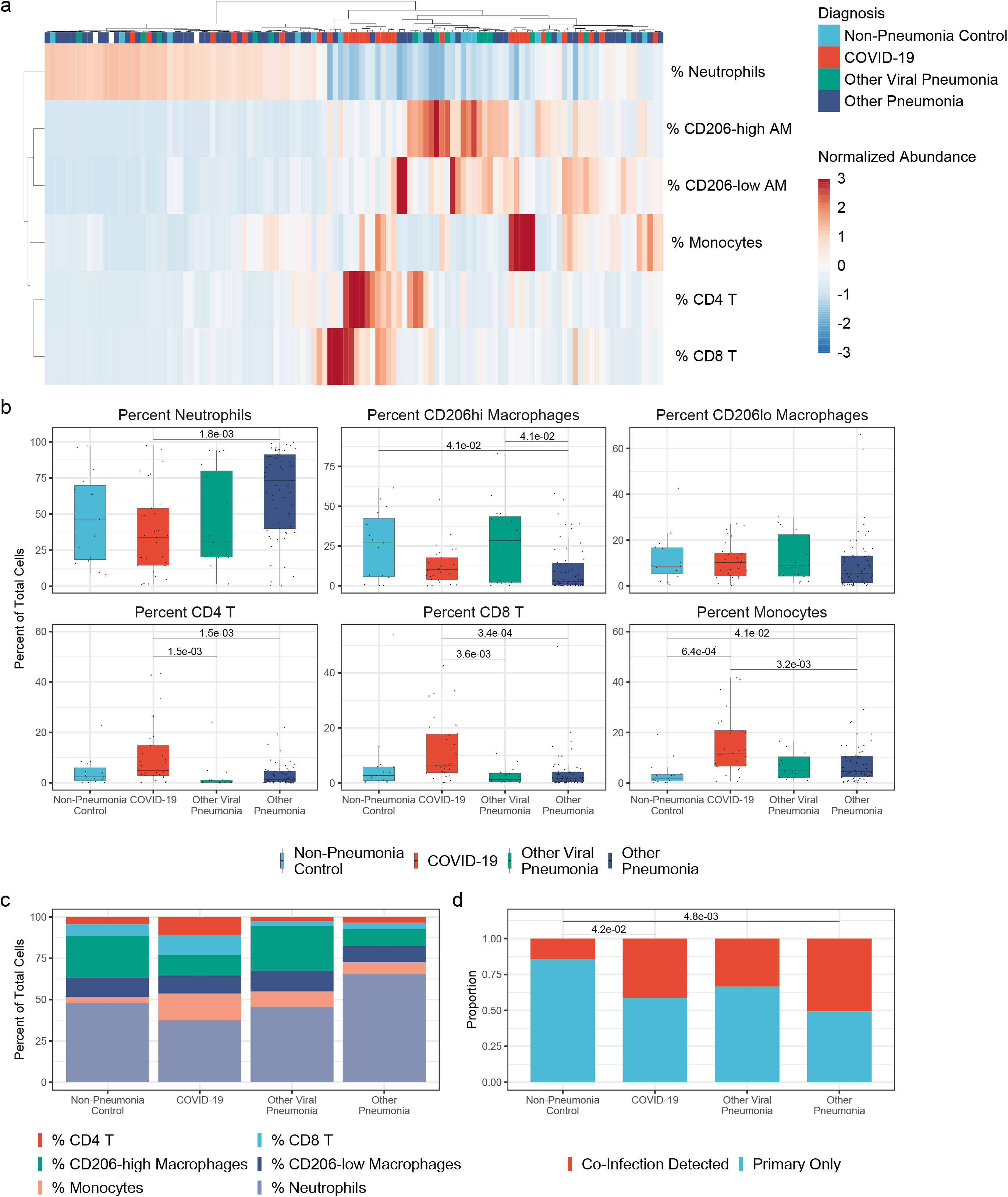
The alveolar space in patients with severe SARS-CoV-2 pneumonia is enriched for T cells and monocytes and lacks neutrophils at the onset of mechanical ventilation. **a**. Hierarchical clustering of BAL samples collected within 48 hours of intubation based on their composition. Samples from patients with non-COVID-19 pneumonia are characterized by neutrophilia while BAL fluid from most patients with SARS-CoV-2 pneumonia is enriched for CD4+ and CD8+ T cells and monocytes. Samples were clustered by Euclidean distance using Ward’s method. **b.** Proportions of cells detected within 48 hours of intubation. Proportion of CD4+ and CD8+ T cells is increased in the COVID-19 cohort (q < 0.05, pairwise Wilcoxon rank-sum tests with FDR correction) and proportion of neutrophils is reduced in these patients, relative to non-viral pneumonia controls (q < 0.05, pairwise Wilcoxon rank-sum tests with FDR correction). Comparisons are not significant unless otherwise noted. **c.** Averaged cell-type compositions in the first 48 hours of intubation, binned by diagnosis. **d.** Comparison of rates of co-infection. Co-infection was defined by detection in a single sample of any bacterial or fungal pathogen listed in **Table 2** by culture or multiplex PCR analysis. No significant differences were observed between the COVID-19 cohort and pneumonia controls (q ≥ 0.05, pairwise Chi-Square tests of proportions with continuity and FDR correction).

### Transcriptomic analysis of flow-sorted alveolar macrophages identifies an interferon response signature as a unique feature of SARS-CoV-2 pneumonia

We performed RNA-Seq on flow-sorted alveolar macrophages from patients with severe SARS-CoV-2 pneumonia (82 samples from 51 patients) and compared them with patients with pneumonia secondary to other causes (29 samples from 23 patients with any other viral pneumonia, 103 samples from 84 patients with non-viral pneumonia), non-pneumonia controls (24 samples from 23 patients) and healthy volunteers (5 samples from 5 volunteers). k-means clustering of the 2,149 most variable genes across diagnoses using a likelihood-ratio test (LRT) identified 5 clusters (**Figure 3a, Supplemental Figure 3a, Supplemental Data Files 1–3**). Notably, patients with COVID-19 clustered together. Cluster 1 contained genes specifically upregulated in patients with COVID-19, relative to patients with other pneumonia and non-pneumonia controls and was characterized by genes involved in the response to interferon (IFN) (GO:0060333 interferon-gamma-mediated signaling pathway; GO:0002474 antigen processing and presentation of peptide antigen via MHC class I; GO:0051607 defense response to virus). Cluster 1 included genes encoding the chemokine *CCL8,* which drives T cell recruitment via CCR5, and *CCL24*, a highly specific T cell chemoattractant (**Figure 3a** and **Supplemental Figure 3b**). Cluster 2 genes were upregulated in patients with early COVID-19 and patients with other pneumonia, but not in non-pneumonia controls and other viral pneumonia. Cluster 2 was characterized by expression of genes associated with monocyte-derived macrophages, such as *CCR2, CCL2, FCN1, F13A1, RNASE1, PLA2G7, FCGR2B, CCL17, CCL7, CCR1, CCR5, MAFB, FCGR2C* (GO:1905517 macrophage migration; GO:0042116 macrophage activation; GO:0006955 immune response). Cluster 3 was characterized by expression of genes involved in monocyte-to-macrophage differentiation, including transcription factors *NR4A1* and *BHLHE40*, *CSF2RA* encoding a subunit of GM-CSF receptor, essential molecular chaperones (*BAG3, DNAJB1, HSPA1B, HSPA1A*), and multiple cytokines and chemokines (*CCL20, IL1A, IL1B, AREG, VEGFA, CXCR4, CXCL3, CXCL5, CXCL1, CXCL8*; GO:0002548 monocyte chemotaxis; GO:0002449 lymphocyte mediated immunity; GO:0032611 interleukin-1 beta production). This cluster was also characterized by unfolded protein response activation resulting from hypoxia, including *HIF1, LAMP3, and DDIT4* (GO:0001666 response to hypoxia). Cluster 3 was enriched for patients with non-viral pneumonia. Genes characterizing cluster 4 were not upregulated in patients with COVID-19 or in healthy controls and included *CD164, MFGE8, IL13RA1,* and *FCGR2A,* suggesting presence of maturing monocyte-derived macrophages. Cluster 5 expressed *FABP4, MSR1, AXL, C1QA, ALOX5, MARCO* and other genes typical for homeostatic resident alveolar macrophages (GO:0006629 lipid metabolic process; GO:0006909 phagocytosis; GO:0043277 apoptotic cell clearance). Interestingly, *IL6*, the protein product of which has been implicated in cytokine storm in severe COVID-19 and serves as a predictor of morbidity and mortality in severe COVID-19 [9], was not among differentially expressed genes and its overall expression was not different between the groups with the exception of healthy controls, where *IL6* transcripts were never detected (**Supplemental Figure 3b**).

**Figure 3.**
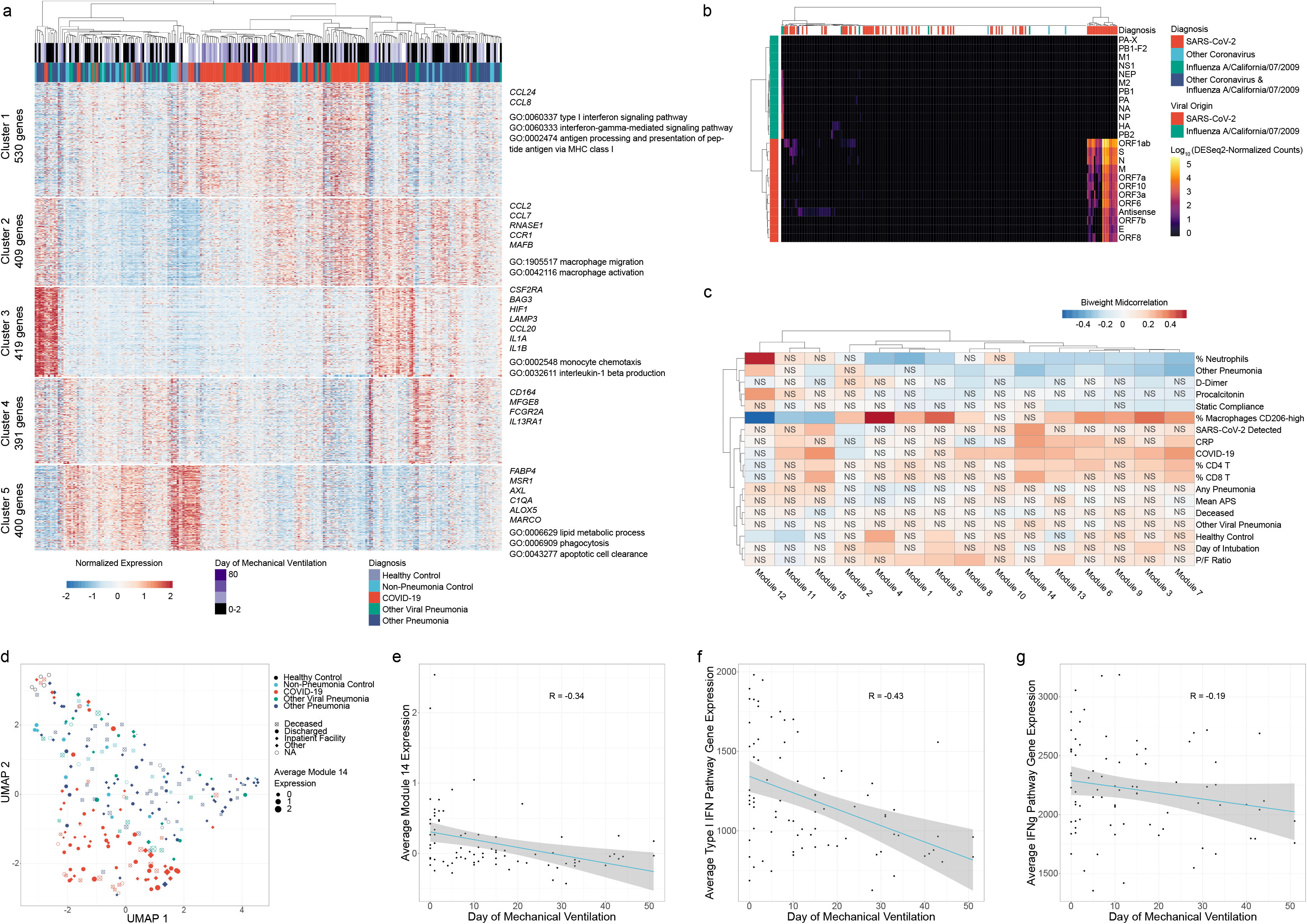
Transcriptomic analysis of alveolar macrophages identifies an interferon response signature as a feature distinguishing SARS-CoV-2 pneumonia from other types of pneumonia. **a**. k-means clustering of the 2,149 most variable genes across diagnoses using a likelihood-ratio test (LRT), columns represent each individual patient and clustered using Ward’s method. Representative genes and biological GO processes are shown for each cluster. Column headers are color-coded by the diagnosis and duration of mechanical ventilation. **b**. Hierarchical clustering of viral reads for SARS-CoV-2 or influenza A/California/07/2009 virus using Ward’s method. Log(DESeq2-normalized counts) are shown. **c**. Weighted correlation network analysis identifies correlation between interferon response genes and SARS-CoV-2 transcripts (module 14) and clinical hallmarks of severe COVID-19: elevated C-reactive protein (CRP) and proportion of CD8 T cells. **d.** UMAP projection of all bulk RNA-seq samples. Average expression of WGCNA module 14 is shown by point diameter. **e–g**. Interferon response signatures in alveolar macrophages from patients with COVID-19 gradually decrease over the course of disease. Correlation between average expression of genes from Module 14 of WGCNA (panel **d**.), GO:0060337 “type I interferon signaling pathway” (panel **e**.), and GO:0060333 “interferon-gamma-mediated signaling pathway” (panel **f**.) and time on mechanical ventilation.

We then asked whether we can detect SARS-CoV-2 transcripts in flow-sorted alveolar macrophages and whether the presence of viral transcripts correlates with specific changes in alveolar macrophage transcriptomes. To allow detection of viral RNA, we aligned sequences to a hybrid genome including human, SARS-CoV-2 and Influenza A/California/07/2009 reference genomes. An additional negative strand transcript, which is transiently formed during viral replication, was added to the SARS-CoV-2 transcriptome to detect replicating virus [10]. We queried macrophage transcriptomes from all patients for the presence of positive- and negative-strand transcripts encoding SARS-CoV-2. We detected SARS-CoV-2 transcripts in macrophage transcriptomes from 67% of patients with PCR-confirmed SARS-CoV-2 infection. In 38% of these patients, we detected both positive- and negative-strand SARS-CoV-2 transcripts (**Figure 3b, Supplemental Figure 3c**). As would be expected during recovery, we also observed a significant negative correlation between abundance of SARS-CoV-2 transcripts in patients with confirmed COVID-19 and days of ventilation (ρ = −0.45, Spearman correlation; **Supplemental Figure 3d**).

In order to identify more specific macrophage gene modules, particularly those that distinguish pneumonia type and outcome, we performed weighted gene coexpression network analysis (WGCNA) (**Figure 3c**, **Supplemental Data File 1**). As predicted, we identified some modules related to composition of the bulk samples, correlating strongly with flow cytometry results. Module 4 exhibited a strong association with the percentage of CD206^hi^ alveolar macrophages, and was characterized by genes expressed in mature tissue-resident alveolar macrophages, including *FABP4* and *PPARG* (R = 0.53) [11]. Module 12 exhibited a strong negative correlation with percentage of CD206^hi^ macrophages, and was identified by expression of genes found in immature monocyte-derived alveolar macrophages including *SPP1* (R = −0.59).

Strikingly, module 14 correlated with the detection of SARS-CoV-2 transcripts, levels of C-reactive protein, CD8+ T cell abundance, and COVID-19 diagnosis (R = 0.30, 0.29, 0.27, 0.26, respectively). Notably, all SARS-CoV-2 genes included in this analysis were assigned to this module, further underscoring disease relevance. In confirmation of the k-means clustering results, this module was highly enriched for type I and type II interferon response genes (GO:0060337 type I interferon signaling pathway; GO:0060333 interferon-gamma-mediated signaling pathway). This association was further confirmed by UMAP projection of all bulk RNA-seq samples, which separated largely by diagnosis with module 14 as a major driver of clustering (**Figure 3d**). Significant negative correlations of average module 14 expression, type I interferon signaling (GO:0060337), and type II interferon signaling (GO:0060333) were observed over the time-course of mechanical ventilation, suggesting reductions during the recovery phase (**Figure 3e–g**).

### Single cell RNA-Seq identifies a positive feedback loop between IFNγ-producing T cells and SARS-CoV-2-infected alveolar macrophages

RNA-Seq analysis of flow-sorted alveolar macrophages suggested that at the time of intubation in patients with severe COVID-19, tissue-resident alveolar macrophages (TRAMs) have been replaced with recruited monocyte-derived alveolar macrophages (MoAM) characterized by an IFN-response signature. To gain insights about early events preceding the onset of COVID-19 respiratory failure we performed single cell RNA-Seq on 5 patients from whom BAL samples were collected within 48 hours of intubation and in which flow cytometry identified distinct populations of CD206^hi^ and CD206^lo^ macrophages. A single patient with bacterial pneumonia was used as a control. Samples were enriched via flow sorting for live cells excluding granulocytes.

Analysis of the integrated COVID-19 dataset resolved clusters corresponding to macrophages, subsets of dendritic cells, epithelial cells (ionocytes, club, ciliated and alveolar epithelial type 1 and type 2 cells), subsets of T cells, B cells and plasma cells (**Figure 4a, Supplemental Data File 4**). Macrophages contained 5 clusters: three subclusters of monocyte-derived alveolar macrophages (all characterized by *CCL2* expression) with different degrees of maturation (MoAM1, *CCL2+IL1R2+CCL8+MRC1-FABP4-*; MoAM2, *CCL2+CCL18+MRC1+FABP4-*; MoAM3, *CCL2+MCEMP1+MRC1+FABP4-*), and two subclusters of resident alveolar macrophages – TRAM1 and TRAM2 both expressing typical TRAM marker genes (*FABP4, RBP4, INHBA*) (**Figure 4b**). All macrophage clusters were represented by cells from 4–5 patients (**Supplemental Figure S4a**).

**Figure 4.**
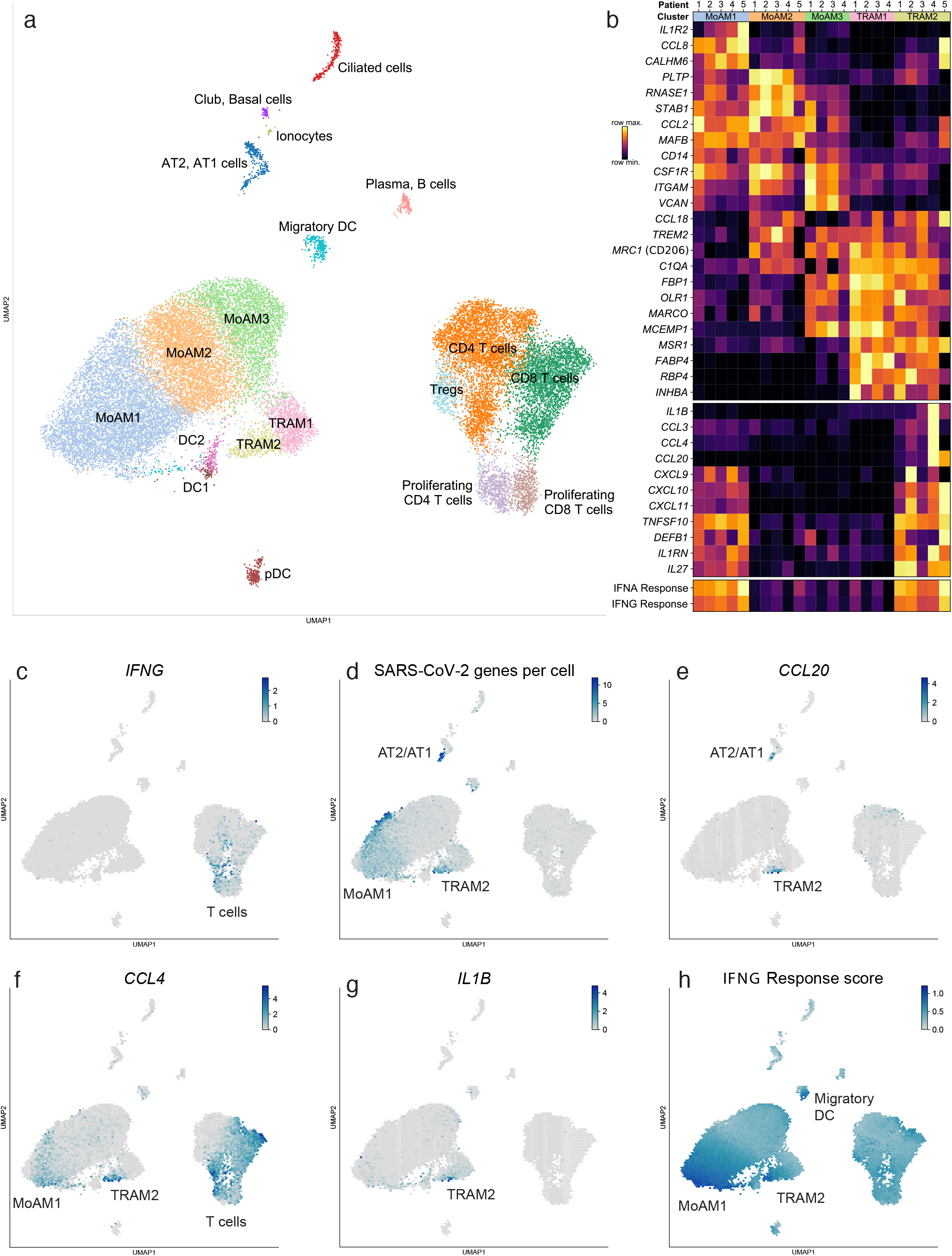
Single cell RNA-Seq identifies a positive feedback loop between IFNγ-producing T cells and SARS-CoV-2-infected alveolar macrophages. **a**. UMAP plot showing integrative analysis of 27,930 cells isolated from 5 patients with severe COVID-19 within 48 hours after intubation. AT1 – alveolar epithelial type 1 cells; AT2 – alveolar epithelial type 2 cells; DC1 – conventional dendritic cells type 1, *CLEC9A*+; DC2 – conventional dendritic cells type 2, *CD1C*+; Migratory DC – migratory dendritic cells, *CCR7*+; pDC – plasmacytoid dendritic cells, *CLEC4C*+; TRAM – tissue-resident alveolar macrophages; MoAM – monocyte-derived alveolar macrophages; Tregs – regulatory T cells, *FOXP3*+. **b.** Heatmap demonstrating expression of the selected genes of interest in two subsets of tissue-resident (TRAM1 and TRAM2) and monocyte-derived (MoAM1, MoAM2, MoAM3) alveolar macrophages. Only 2 cells from patient #5 belonged to cluster MoAM3 and hence were excluded. **c**. Expression of *IFNG* is restricted to T cells. **c.** Detection of SARS-CoV-2 transcripts. Plot shows cumulative number of viral genes plus negative strand SARS-CoV-2. **e-g**. Specific upregulation of selected cytokines and chemokines in TRAM2: *CCL20* (panel **e**), *CCL4* (panel **f**), *IL1B* (panel **g**). Density projection plots, expression averaged within hexagonal areas on UMAP. **h**. Increased expression of IFNγ-responsive genes in MoAM1 and TRAM2 clusters characterized by the presence of SARS-CoV-2 transcripts.

As our analysis of transcriptomic data from alveolar macrophages suggested that SARS-CoV-2 pneumonia is uniquely associated with the activation of pathways induced by interferons, we looked for the expression of type I interferons in our single cell dataset. We did not detect type I interferon expression in any of the sequenced cells (data not shown). Similarly, we did not detect type I interferon transcripts in other publicly available single cell RNA-Seq datasets obtained from BAL fluid [12,13]. In contrast, expression of type II interferon – *IFNG* – was detected in T cells from all 5 patients (**Figure 4c, Supplemental Figure 4b**). These results suggest the interferon response gene signature we observed in flow-sorted alveolar macrophages is induced by IFNγ released from activated T cells.

We next looked for SARS-CoV-2 positive- and negative-strand transcripts within our data. As expected, both were detected in epithelial cells and migratory *CCR7*+ dendritic cells. Interestingly, they were also detected in MoAM1 and TRAM2 subclusters (**Figure 4d, Supplemental Figure 4c**). Coronaviruses generate large numbers of positive strand transcripts from a single negative strand transcript [10,14]. Consistent with this known biology, we detected more transcripts for positive compared with negative strands in both our single cell and bulk RNA-Seq data **Supplementary Figure 4d**). These data suggest that alveolar macrophages harbor virus and suggest they may support viral replication as has been reported for SARS-CoV and MERS-CoV [15–17].

Interestingly, TRAM2 harboring SARS-CoV-2 showed distinct clustering compared with uninfected TRAM1 (clustering was performed without SARS-CoV-2 transcripts). Importantly, these TRAM2 also clustered differently when compared with TRAM from a patient with bacterial pneumonia (**Supplemental Figure 4e, f, Supplemental Data File 5**). Marker genes distinguishing infected TRAM2 from non-infected TRAM1 included several chemokines and cytokines important for T cell recruitment. These include *CCL20*, which attracts dendritic cells and T cells via CCR6 and only weakly attracts neutrophils (**Figure 4b, e**)*, CXCL10* and *CXCL11*, which attract T cells and plasmacytoid dendritic cells via CXCR3, and *CCL4*, which attracts monocytes, NK and T cells via CCR5 (**Figure 4b, f**). These cells also expressed genes important in the innate immune response to virus. These include *IL1B* (**Figure 4b, g**), *TNFSF10*, a member of TNF ligand family, and the antimicrobial peptide defensin B (*DEFB1*) (**Figure 4b, Supplemental Data File 6**). Finally, these cells were marked by increased expression of interferon response genes (*ISG15, IFIT1, IFIT2, IFIT3, MX1* and others). Consistent with this observation, we found higher average levels of expression of a curated list of IFN-response genes in infected (TRAM2) compared with non-infected (TRAM1) alveolar macrophages (**Figure 4b,h; Supplemental Figure 4g**). These results indicate a positive feedback loop in which SARS-CoV-2-infected alveolar macrophages release chemokines that recruit T cells, which release IFNγ to further activate the infected alveolar macrophages.

High serum levels of IL-6 and its transcriptional target C-reactive protein have been observed in patients with severe COVID-19. IL-6 induces the transcription of clotting factors in the liver and tissue factor in the endothelium that promote thrombosis [18]. These mechanisms are thought to contribute to the microvascular thrombosis in the lung and other tissues observed in autopsy studies conducted on patients with COVID-19, which in turn is thought to contribute to multiple organ dysfunction [19,20]. As increased transcripts encoding IL-6 have not been observed in single cell RNA-Seq studies of the peripheral blood [21], some have suggested IL-6 is generated by inflammatory cells in the alveolus [20]. The overall expression of *IL6* was low, and was not restricted to a specific cell type in our single cell RNA-Seq dataset. Despite this, the levels of C-reactive protein at the time of intubation were increased in patients with severe COVID-19, relative to the upper limit of the normal range of 10 mg/L (P = 2.5×10^−3^, one-sample Wilcoxon rank-sum test; **Supplemental Figure S1c**). These results suggest that the resident and recruited immune cells in the infected alveoli do not directly contribute to systemic increase of IL-6 (**Supplemental Figure S4h**).

### Severe SARS-CoV-2 pneumonia unfolds slowly over time

Our studies of BAL fluid collected from patients with COVID-19 within 48 hours of intubation revealed an already established feedback loop between activated T cells and infected alveolar macrophages. This stable, slowly amplifying feedback loop is consistent with the long duration between symptom onset and the development of respiratory failure in patients with COVID-19 (5–9 days) and the prolonged course of mechanical ventilation in many of these patients (**Figure 1a, h, i)** [1]. A prediction from this model is that these circuits will persist over time until the virus is cleared. Hence, we examined serial BAL fluid samples collected from patients with severe SARS-CoV-2 pneumonia over time (**Figure 5a, b**). We used clinical measures performed on the BAL fluid (quantitative culture or multiplex PCR for respiratory pathogens) to identify bacterial superinfections, defined as detection of a new bacterial pathogen acquired during the hospitalization. In the absence of a detected bacterial superinfection, BAL fluid remained non-neutrophilic in 62% of samples from patients admitted with SARS-CoV-2 pneumonia. BAL neutrophilia developed in 50% of patients with COVID-19 who developed bacterial superinfection. Hierarchical clustering of the composition of BAL samples from all time points demonstrated that in comparison to samples from other types of pneumonia samples from patients with COVID-19 were enriched for T cells irrespective of the time of BAL collection (**Figure 5a, b**) or presence or absence of superinfection (**Figure 5c**).

**Figure 5.**
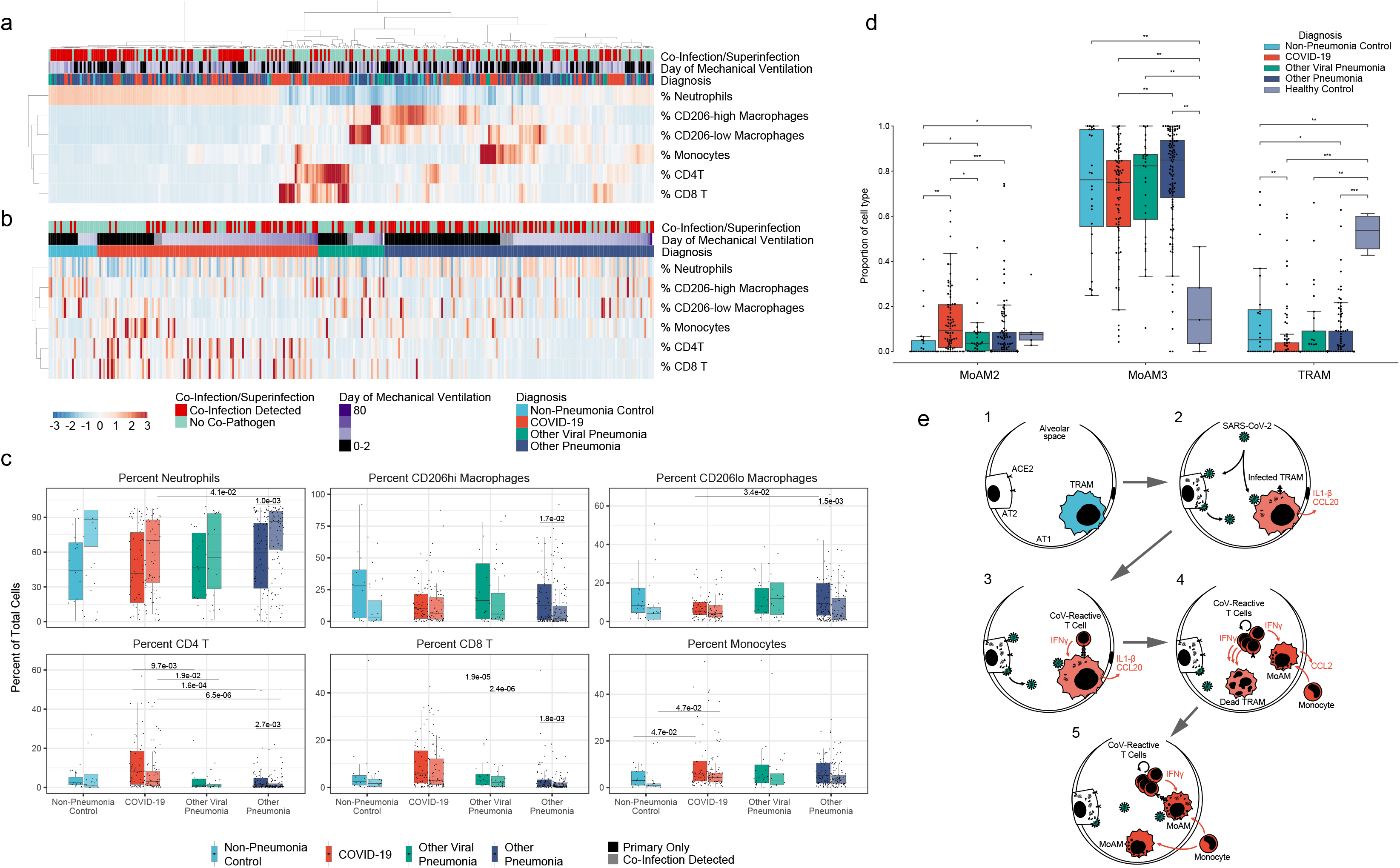
The BAL fluid from patients with SARS-CoV-2 pneumonia is dominated by T cells during the course of the disease irrespective of superinfection status. **a**. Hierarchical clustering of BAL samples based on their composition demonstrates several clusters of patients with COVID-19 dominated by T cells, monocytes and macrophages. Samples were clustered by Euclidean distance using Ward’s method. **b.** Clustering of BAL samples from *panel a*, grouped by diagnosis and ordered by the duration of mechanical ventilation. Samples from patients with COVID-19 were dominated by T cells, monocytes and macrophages. **c.** Proportions of cells across all diagnosis between patients without and with superinfection. A substantial enrichment of CD4+ and CD8+ T cells was observed in the COVID-19 cohort in comparison to non-viral pneumonia irrespective of whether a secondary infection was detected (q < 0.05, pairwise Wilcoxon rank-sum tests with FDR correction). **d**. Ongoing recruitment of more immature MoAM2, relative decrease of more mature MoAM3 and loss of TRAMs in patients with COVID-19 in comparison to other types of pneumonia. Deconvolution of bulk RNA-seq gene expression profiles (**Figure 3a**) was performed using signatures from integrated single-cell RNA-seq (**Figure 4a**). Asterisks indicate statistical significance, Wilcoxon rank-sum tests with FDR correction: * p < 0.05, ** p < 0.01, *** p < 0.001. **e**. Schematic illustrating interpretation of the main findings. 1. Normal alveolus contains *ACE2*-expressing alveolar type 1 and type 2 cells (AT1 and AT2, correspondingly) and tissue-resident alveolar macrophages (TRAM). 2. SARS-CoV-2 infects AT2 cells and TRAM. Infected TRAM are activated and express *IL1B* and T cell chemokines. 3. Cross-reactive or *de novo* generated T cells recognize SARS-CoV-2 antigens presented by TRAM and produce IFNγ, further activating TRAM to produce cytokines and chemokines. 4. Activated T cells proliferate and continue to produce IFNγ, eventually leading to death of infected TRAM and recruitment of monocytes, which rapidly differentiate into monocyte-derived alveolar macrophages (MoAM). MoAM in turn produce chemokines that further promote T cell and monocyte recruitment, thus sustaining pneumonia. 5. Recruited MoAM either become mature and repopulate alveoli or become infected with SARS-CoV-2, continuing to present antigens to T cells and maintain the feedback loop.

Our data suggest that composition of immune cells in SARS-CoV-2 pneumonia is relatively stable and continues to unfold slowly over time, resulting in a loss of tissue-resident alveolar macrophages and continuous recruitment of monocyte-derived alveolar macrophages. To test this hypothesis, we performed cell type deconvolution to estimate the proportion of individual cell types in bulk RNA-seq data from flow-sorted alveolar macrophages using cell type-specific signatures from single cell RNA-seq. This approach confirmed that only a small portion of the samples from patients with pneumonia contained tissue-resident alveolar macrophages. Instead, the majority of alveolar macrophages were mature monocyte-derived alveolar macrophages (MoAM3) (**Figure 5d, Supplemental Figure 4i**). Interestingly, samples from patients with COVID-19 contained more immature MoAM2 alveolar macrophages than samples from patients with other types of pneumonia, supporting our hypothesis that SARS-CoV-2 pneumonia is characterized by the ongoing recruitment of monocyte-derived alveolar macrophages.

## Discussion

Our data represent the earliest sampling of the alveolar space of patients with COVID-19 pneumonia reported to date [12,22]. It is possible to use these data to speculate about early events that lead to pneumonia in patients with SARS-CoV-2 infection (**Figure 5e**). Strong evidence suggests SARS-CoV-2 initially infects and replicates in epithelial cells in the nasopharynx, which express relatively high levels of *ACE2* in comparison with epithelial cells in airways or the distal lung [23,24]. There the virus replicates and escapes clearance by suppressing type I interferon responses [25–27] and broadly disrupting protein translation [28]. Whether by progressive movement distally in the tracheobronchial tree or via aspiration of nasopharyngeal contents, some virus gains access to the distal alveolar space. In the alveolar space, we confirm that SARS-CoV-2 infects alveolar epithelial cells and alveolar macrophages [27]. Alveolar macrophages might be infected with SARS-CoV-2 by 1) phagocytosis of infected epithelial cells, 2) via direct infection, as was shown for SARS-CoV and MERS-CoV [15,16] or 3) through other mechanisms, such as antibody mediated enhancement as was shown for SARS-CoV [17,29]. Indeed, it is estimated that ~90% of adults have antibodies against one of four major seasonal coronaviruses [30], and antibodies cross-reacting with SARS-CoV-2 might facilitate viral entry into alveolar macrophages.

While tissue-resident alveolar macrophages are poor antigen-presenting cells and do not migrate to lymph nodes to participate in conversion of naive T cells into effector T cells [31], a low level of antigen presentation by alveolar macrophages in the alveolar space might be sufficient to drive activation of pre-existing memory T cells that target endemic seasonal coronaviruses, but cross-react with SARS-CoV-2. Existence of such cross-reactive memory T cells has been reported for SARS-CoV [32] and, recently, SARS-CoV-2 [33–36]. This mechanism might explain the epidemiology of SARS-CoV-2, which disproportionately affects elderly individuals, while children and young adults often have only mild symptoms, despite having viral load in upper airways compatible to adults [37,38]. Children and young individuals have fewer encounters with seasonal coronaviruses during their lifetime than elderly individuals and therefore would have fewer cross-reactive antibodies or memory T cells.

We did not detect substantial expression of type I interferons in our datasets. This could be either related to the timing of sampling, undersampling cell types producing type I interferons, or inhibition of type I interferon production by SARS-CoV-2 proteins [25,39]. However, we readily detected IFNγ production by T cells. This supports a model where infection and activation of tissue-resident alveolar macrophages with SARS-CoV-2, followed by antigen presentation to cross-reactive or SARS-CoV-2-specific T cells, leads to a delayed interferon response. This in turn amplifies the inflammatory response by recruiting proinflammatory monocyte-derived macrophages, as has been suggested from animal models of SARS-CoV infection [40]. Interestingly, in agreement with our observation that the inflammatory environment in the lung during severe COVID-19 is characterized by the paucity of neutrophils, we found that alveolar macrophages produced chemokines preferentially driving recruitment of T cells and monocytes, but not neutrophils.

Our results also explain the slow progression of SARS-CoV-2 pneumonia relative to other respiratory viruses, most notably influenza A. Specifically, the time from the onset of symptoms to respiratory failure in patients with SARS-CoV-2 infection is 5–9 days, compared with 1–3 days or even less in patients with influenza A infection[4]. Consistent with a slower course, the duration of illness is longer in patients with severe SARS-CoV-2 infection. While both viruses might gain access to the distal lung by similar mechanisms, the sialic acid residues that serve as receptors for influenza A are abundantly expressed in alveolar type 2 cells [23]. In contrast, single cell RNA-Seq atlases of the human lung and sm-FISH studies of the normal lung show that only a small number of alveolar epithelial cells express *ACE2* [23,24]. Thus, while influenza A would infect large numbers of cells causing widespread injury, rapid viral replication and robust antiviral responses, infection by SARS-CoV-2 is likely to lead to more localized areas of infection. These more localized areas of infection and injury are consistent with radiographic patterns of early COVID-19, and the presence of discrete radiographic abnormalities in asymptomatic COVID-19 patients [41]. Because tissue-resident alveolar macrophages have limited motility [42], the uptake and persistence of SARS-CoV-2-infected alveolar macrophages would not be predicted to spread the virus to adjacent lung regions. However, recruited monocyte-derived alveolar macrophages, which we found can also harbor replicating SARS-CoV-2, are typically more numerous, and could conceivably spread the virus to adjacent alveoli. In each new area of infection, positive feedback loops between alveolar macrophages harboring the virus and activated T cells could promote ongoing injury and systemic inflammation. This model would also explain the heterogeneity we observed in alveolar macrophages recovered from the same individual. Different stages of SARS-CoV-2 infection might be spatially localized to subsegmental regions such that the BAL procedure samples alveoli in various stages of infection. Further studies using spatial techniques and a time-course analysis of patients with early disease will address these questions.

## Acknowledgments and funding

Northwestern University Flow Cytometry Core Facility is supported by NCI Cancer Center Support Grant P30 CA060553 awarded to the Robert H. Lurie Comprehensive Cancer Center. Cell sorting was performed on BD FACSAria SORP cell sorter purchased through the support of NIH 1S10OD011996-01.

This research was supported in part through the computational resources and staff contributions provided by the Genomics Compute Cluster which is jointly supported by the Feinberg School of Medicine, the Center for Genetic Medicine, and Feinberg's Department of Biochemistry and Molecular Genetics, the Office of the Provost, the Office for Research, and Northwestern Information Technology. The Genomics Compute Cluster is part of Quest, Northwestern University's high performance computing facility, with the purpose to advance research in genomics and world peace.

Rogan Grant was supported by NIH grant T32AG020506-18.

Luisa Morales-Nebreda was supported by T32HL076139 and F32HL151127.

GR Scott Budinger was supported by NIH grants U19AI135964, P01AG049665, R01HL147575 and Veterans Affairs grant I01CX001777.

Alexander V. Misharin was supported by NIH grants U19AI135964, P01AG049665, R56HL135124, R01HL153312 and NUCATS COVID-19 Rapid Response Grant.

Benjamin D. Singer was supported by NIH awards K08HL128867, U19AI135964, R01HL149883, and P01AG049665.

Richard G. Wudnerink was supported by NIH grant U19AI135964 and a GlaxoSmithKline Distinguished Scholar in Respiratory Health grant from the CHEST Foundation.

## Consortia: The NU SCRIPT Study Investigators

A. Christine Argento, Ajay A. Wagh, Alan R. Hauser, Alexander V. Misharin, Alexis Rose Wolfe, Ali Shilatifard, Alvaro Donayre, Anjali Thakrar, Ankit Bharat, Ann A. Wang, Anna E. Pawlowski, Anne R. Levenson, Anthony M. Joudi, Benjamin D. Singer, Betty Tran, Cara J. Gottardi, Catherine A. Gao, Chao Qi, Chiagozie O. Pickens, Chitaru Kurihara, Clara J Schroedl, Curt M. Horvath, Daniel Meza, Daniel Schneider, Darryl A. Abbott, David D. Odell, David W. Kamp, Deborah R. Winter, Egon A. Ozer, Elizabeth T Bartom, Elizabeth S. Malsin, Emily J. Rendleman, Emily M. Leibenguth, Estefany R. Guzman, Firas Wehbe, Gabrielle Y. Liu, Gaurav T. Gadhvi, GR Scott Budinger, Helen K. Donnelly, Heliodoro Tejedor Navarro, Hermon Kihshen, Hiam Abdala-Valencia, Isaac A. Goldberg, Jacob I. Sznajder, Jacqueline M. Kruser, James M. Walter, Jane E. Dematte, Jasmine Le, Jason M. Arnold, Joanne C. Du, John Coleman, Joseph Isaac Bailey, Joseph S. Deters, Justin A. Fiala, Justin Starren, Karen M. Ridge, Katharine Secunda, Kathleen Aren, Khalilah L. Gates, Kristy Todd, Lango Sichizya, Lindsey D. Gradone, Lindsey N. Textor, Lisa F. Wolfe, Lorenzo L. Pesce, Luisa Morales-Nebreda, Luís A. Nunes Amaral, Madeline L Rosenbaum, Manoj Kandpal, Manu Jain, Marc A. Sala, Mark Saine, Mary Carns, Melissa Querrey, Mengjia Kang, Michael J. Alexander, Michael J. Cuttica, Michelle Hinsch Prickett, Nicholas D. Soulakis, Nicole Borkowski, Nikolay S. Markov, Orlyn R. Rivas, Paul A. Reyfman, Pearl D. Go, Peter H. S. Sporn, Phillip R. Cooper, Prasanth Nannapaneni, Rade Tomic, Radhika Patel, Rafael Garza-Castillon, Ravi Kalhan, Richard G. Wunderink, Rogan A. Grant, Ruben J. Mylvaganam, Sean Smith, Patrick C. Seed, Samuel S. Kim, Samuel W.M. Gatesy, Sanket Thakkar, Sarah Ben Maamar, SeungHye Han, Sharon R. Rosenberg, Sophia Nozick, Stefan J. Green, Suchitra Swaminathan, Susan R. Russell, Taylor A. Poor, Theresa A. Lombardo, Thomas Stoeger, Todd Shamaly, Yuliya Politanska, Zasu M. Klug, Ziyan Lu, Ziyou Ren.

## Methods

### Human subjects

All human subjects research was approved by the Northwestern University Institutional Review Board. Samples from patients with COVID-19, viral pneumonia, other pneumonia and non-pneumonia controls were collected from participants enrolled in Successful Clinical Response In Pneumonia Therapy (SCRIPT) study STU00204868. Alveolar macrophages from healthy volunteers were obtained under study STU00206783. All subjects or their surrogates provided informed consent.

Patients ≥ 18 years of age with suspicion of pneumonia based on clinical criteria (including but not limited to fever, radiographic infiltrate, and respiratory secretions) were screened for enrollment into the SCRIPT study. Inability to safely obtain BAL or NBBAL were considered exclusion criteria. In our center, patients with respiratory failure are intubated based on the judgement of bedside clinicians for worsening hypoxemia, hypercapnia, or work of breathing refractory to high-flow oxygen or non-invasive ventilation modes. Extubation occurs based on the judgement of bedside clinicians following a trial of spontaneous breathing in patients demonstrating physiologic improvement in their cardiorespiratory status during their period of mechanical ventilation.

Management of patients with COVID-19 was guided by protocols published and updated on the Northwestern Medicine website as new information became available over the pandemic. Clinical laboratory testing including studies ordered on bronchoalveolar lavage fluid was at the discretion of the care team, however, quantitative cultures, multiplex PCR (BioFire Film Array Respiratory 2 panel), and automated cell count and differential were recommended by local ICU protocols. Most patients also underwent urinary antigen testing for *Streptococcus pneumoniae* and *Legionella pneumophilia* on admission. Clinicians were encouraged to manage all patients, including those with COVID-19, according to ARDSnet guidelines including the use of a higher PEEP/lower FIO2 strategy for those with severe hypoxemia. Prone positioning (16 hours per day) was performed in all patients with a PaO2/FiO2 <150 who did not have contraindications. In those who had a response to prone positioning evident by improved oxygenation, prone positioning was repeated. Esophageal balloon catheters (Cooper Surgical) were placed at the discretion of the care team to estimate transpulmonary pressure and optimize PEEP, particularly in patients with a higher than normal BMI.

Pneumonia category adjudication was performed by five critical care physicians (RGW, JMK, BDS, COP, JMW) using a standardized reporting tool (Supplemental Data File 7). Clinical laboratory data were obtained from the Northwestern Medicine Enterprise Data Warehouse using Structured Query Language (SQL). APS and SOFA scores were generated from the Electronic Health Record using previously validated programming.

### NBBAL and BAL procedures

Consent was obtained from patients or legal decision makers for the bronchoscopic procedures. Bronchoscopic BAL was performed in intubated ICU patients with flexible, single-use Ambu aScope (Ambu) devices. Patients were given sedation and topical anesthetic at the physician proceduralist’s discretion. Vital signs were monitored continuously throughout the procedure. The bronchoscope was wedged in the segment of interest based on available chest imaging or intra-procedure observations, aliquots of 30 cc of normal saline at a time, generally 90–120 cc total, were instilled and aspirated back. The fluid returned following the first aliquot was routinely discarded. Samples were split (if sufficient return volume was available) and sent for clinical studies and an aliquot reserved for research. A similar procedure was applied to non-bronchoscopic BAL (NBBAL); however NBBAL was performed with directional but not visual guidance, and as usual procedural care by a respiratory therapist rather than a pulmonologist[6].

For the bronchoscopies performed in the COVID-19 patients, additional precautions were taken to minimize the risk to healthcare workers including only having essential providers present in the room, clamping of the endotracheal tube, transient disconnection of the inspiratory limb from the ventilator, and preloading of the bronchoscope through the adapter; sedation and neuromuscular blockade to prevent cough, was administered for these procedures at the physician’s discretion. In most cases the early bronchoscopy was performed immediately after intubation.

### Flow cytometry and cell sorting

NBBAL and BAL samples were filtered through a 70 um cell strainer, pelleted by centrifugation at 300 rcf for 10 min at 4C, followed by hypotonic lysis of red blood cells with 2 ml of BD PharmLyse reagent for 2 min. Lysis was stopped by adding 18 ml of MACS buffer. Cells were pelleted again and resuspended in 100 ul of Fc-Block (Human TruStain FcX, Biolegend), and 10 ul aliquot was taken for counting using K2 Cellometer (Nexcelom) with AO/PI reagent. The volume of Fc-Block was adjusted so the concentration of cells was always less than 5×10^7^ cells/ml and the fluorophore-conjugated antibody cocktail was added in 1:1 ratio (**Table 3**). After incubation at 4C for 30 min cells were washed with 5 ml of MACS buffer, pelleted by centrifugation and resuspended in 500 ul of MACS buffer + 2 ul of SYTOX Green viability dye (ThermoFisher). Cells were sorted on FACS Aria II SORP instrument using 100 um nozzle and 20 psi pressure. Cells were sorted into 300 ul of MACS buffer for bulk RNA-seq or 300 ul of 2% BSA in PBS for single cell RNA-Seq. Sample processing was performed in BSL-2 facility using BSL-3 practices.

**Table 3:**
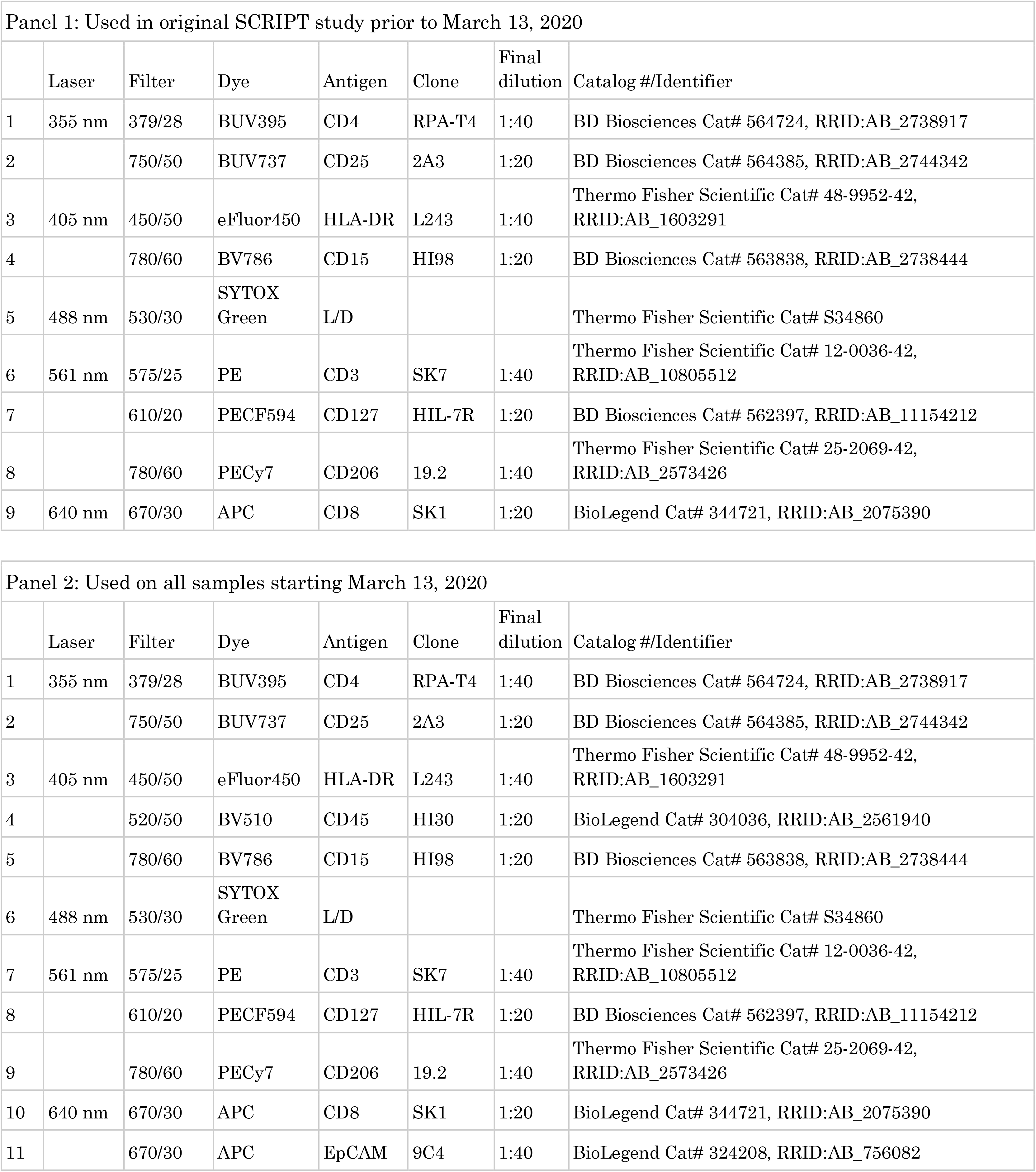
Flow cytometry panels used in this study.

| Panel 1: Used in original SCRIPT study prior to March 13, 2020 |  |  |  |  |  |  |  |
| --- | --- | --- | --- | --- | --- | --- | --- |
|  | Laser | Filter | Dye | Antigen | Clone | Final dilution | Catalog #/Identifier |
| 1 | 355 nm | 379/28 | BUV395 | CD4 | RPA-T4 | 1:40 | BD Biosciences Cat# 564724, RRID:AB_2738917 |
| 2 |  | 750/50 | BUV737 | CD25 | 2A3 | 1:20 | BD Biosciences Cat# 564385, RRID:AB_2744342 |
| 3 | 405 nm | 450/50 | eFluor450 | HLA-DR | L243 | 1:40 | Thermo Fisher Scientific Cat# 48-9952-42, RRID:AB_1603291 |
| 4 |  | 780/60 | BV786 | CD15 | HI98 | 1:20 | BD Biosciences Cat# 563838, RRID:AB_2738444 |
| 5 | 488 nm | 530/30 | SYTOX Green | L/D |  |  | Thermo Fisher Scientific Cat# S34860 |
| 6 | 561 nm | 575/25 | PE | CD3 | SK7 | 1:40 | Thermo Fisher Scientific Cat# 12-0036-42, RRID:AB_10805512 |
| 7 |  | 610/20 | PECF594 | CD127 | HIL-7R | 1:20 | BD Biosciences Cat# 562397, RRID:AB_11154212 |
| 8 |  | 780/60 | PECy7 | CD206 | 19.2 | 1:40 | Thermo Fisher Scientific Cat# 25-2069-42, RRID:AB_2573426 |
| 9 | 640 nm | 670/30 | APC | CD8 | SK1 | 1:20 | BioLegend Cat# 344721, RRID:AB_2075390 |

| Panel 2: Used on all samples starting March 13, 2020 |  |  |  |  |  |  |  |
| --- | --- | --- | --- | --- | --- | --- | --- |
|  | Laser | Filter | Dye | Antigen | Clone | Final dilution | Catalog #/Identifier |
| 1 | 355 nm | 379/28 | BUV395 | CD4 | RPA-T4 | 1:40 | BD Biosciences Cat# 564724, RRID:AB_2738917 |
| 2 |  | 750/50 | BUV737 | CD25 | 2A3 | 1:20 | BD Biosciences Cat# 564385, RRID:AB_2744342 |
| 3 | 405 nm | 450/50 | eFluor450 | HLA-DR | L243 | 1:40 | Thermo Fisher Scientific Cat# 48-9952-42, RRID:AB_1603291 |
| 4 |  | 520/50 | BV510 | CD45 | HI30 | 1:20 | BioLegend Cat# 304036, RRID:AB_2561940 |
| 5 |  | 780/60 | BV786 | CD15 | HI98 | 1:20 | BD Biosciences Cat# 563838, RRID:AB_2738444 |
| 6 | 488 nm | 530/30 | SYTOX Green | L/D |  |  | Thermo Fisher Scientific Cat# S34860 |
| 7 | 561 nm | 575/25 | PE | CD3 | SK7 | 1:40 | Thermo Fisher Scientific Cat# 12-0036-42, RRID:AB_10805512 |
| 8 |  | 610/20 | PECF594 | CD127 | HIL-7R | 1:20 | BD Biosciences Cat# 562397, RRID:AB_11154212 |
| 9 |  | 780/60 | PECy7 | CD206 | 19.2 | 1:40 | Thermo Fisher Scientific Cat# 25-2069-42, RRID:AB_2573426 |
| 10 | 640 nm | 670/30 | APC | CD8 | SK1 | 1:20 | BioLegend Cat# 344721, RRID:AB_2075390 |
| 11 |  | 670/30 | APC | EpCAM | 9C4 | 1:40 | BioLegend Cat# 324208, RRID:AB_756082 |

### Bulk RNA-Seq of flow-sorted alveolar macrophages

Immediately after sorting, cells were pelleted by centrifugation and lysed in 350 ul of RLT Plus lysis buffer (Qiagen) supplemented with 2-mercaptoethanol. Lysed cells were stored at −80C until RNA isolation using AllPrep DNA/RNA Micro kit according to manufacturer’s protocol (Qiagen). RNA quality and quantity were assessed using TapeStation 4200 High Sensitivity RNA tapes (Agilent) and RNA-Seq libraries were prepared from 250 pg of total RNA using SMARTer Stranded Total RNA-Seq Kit v2 (Takara Bio). After QC using TapeStation 4200 High Sensitivity DNA tapes (Agilent) dual indexed libraries were pooled and sequenced on a NextSeq 500 instrument (Illumina), 75 cycles, single-end, to an average sequencing depth of 19.55M reads.

FASTQ files were generated using bcl2fastq (Illumina). To enable detection of viral RNA, a custom hybrid genome was prepared by joining FASTA, GFF, and GTF files for GRCh37.87, SARS-CoV-2 (NC_045512.2) – virus causing COVID-19, and Influenza A/California/07/2009 (GCF_001343785.1), which was the dominant strain of influenza throughout BAL fluid collection at our hospital [43]. An additional negative strand transcript spanning the entirety of the SARS-CoV-2 genome was then added to the GTF and GFF files to enable detection of SARS-CoV-2 replication. Normalized counts tables later revealed extremely high enrichment of SARS-CoV-2 transcripts in diagnosed COVID-19 patients, and strong enrichment of IAV genes in patients marked as other viral pneumonia. To facilitate reproducible analysis, samples were processed using the publicly available nf-core/RNA-seq pipeline version 1.4.2 implemented in Nextflow 19.10.0 using Singularity 3.2.1-1 with the minimal command `nextflow run nf-core/rnaseq −r 1.4.2 – singleEnd −profile singularity -unStranded --three_prime_clip_r2 3` [44–46]. Briefly, lane-level reads were trimmed using trimGalore! 0.6.4 and aligned to the hybrid genome described above using STAR 2.6.1d [47]. Gene-level assignment was then performed using featureCounts 1.6.4 [48].

#### Bulk differential expression analysis (DEA)

All analysis was performed using custom scripts in R version 3.6.3 using the DESeq2 version 1.26.0 framework[49]. Correspondence between lanes was first confirmed by principal component analysis before merging counts using the command `collapseReplicates()`. One outlier sample with low RIN score and exhibiting extreme deviation on PCA and poor alignment and assignment metrics was excluded from downstream analysis. For differential expression analysis (DEA), both MoAM content from flow cytometry data and diagnosis were used as explanatory factors. A “local” model of gene dispersion was employed as this better fit dispersion trends without obvious overfitting; otherwise default settings were used.

#### K-means clustering of bulk samples

For k-means clustering, a custom-built function was used (available at https://github.com/NUPulmonary/utils/blob/master/R/k_means_figure.R). Briefly, variable genes were identified using a likelihood-ratio test (LRT) with local estimates of gene dispersion in DESeq2 with diagnosis as the “full” model, and a reduced model corresponding to intercept, alone. Genes with q ≥ 0.05 were discarded. Extant genes were then clustered using the Hartigan-Wong method with 25 random sets and a maximum of 1000 iterations using the kmeans function in R stats 3.6.3. Samples were then clustered using Ward’s method and plotted using pheatmap version 1.0.12. GO term enrichment was then determined using Fisher’s exact test in topGO version 2.38.1, with org.Hs.eg.db version 3.10.0 as a reference.

#### Weighted gene coexpression network analysis (WGCNA)

WGCNA was performed manually using WGCNA version 1.69 with default settings unless otherwise noted [50]. Genes with counts >5 and detection in at least 10% of samples were included in analysis. To best capture patterns of co-regulation, a signed network was used. Using the pickSoftThreshold function, we empirically determined a soft threshold of 7 to best fit the network structure. A minimum module size of 30 was chosen to isolate relatively large gene modules. Module eigengenes were then related back to patient and sample metadata using biweight midcorrelation. Module GO enrichment was then determined as above using Fisher’s exact test in topGO version 2.38.1, with org.Hs.eg.db version 3.10.0 as a reference. UMAP plotting was performed using uwot version 0.1.8 using the first 20 principal components of the same genes used in WGCNA analysis after Z-scaling and centering, with a minimum distance of 0.2 [51]. Default parameters were otherwise used.

### Single cell RNA-Seq of flow-sorted BAL cells

Cells were sorted into 2% BSA in DPBS, pelleted by centrifugation at 300 rcf for 5 min at 4C, resuspended in 0.1% BSA in DPBS to ~1000 cells/ul concentration. Concentration was confirmed using K2 Cellometer (Nexcelom) with AO/PI reagent and ~5,000–10,000 cells were loaded on 10x Genomics Chip A with Chromium Single Cell 5’ gel beads and reagents (10x Genomics). Libraries were prepared according to manufacturer protocol (10x Genomics, CG000086_RevM). After quality check single cell RNA-seq libraries were pooled and sequenced on NovaSeq 6000 instrument.

Data was processed using Cell Ranger 3.1.0 pipeline (10x Genomics). To enable detection of viral RNA, reads were aligned to a custom hybrid genome containing GRCh38.93 and SARS-CoV-2 (NC_045512.2). An additional negative strand transcript spanning the entirety of the SARS-CoV-2 genome was then added to the GTF and GFF files to enable detection of SARS-CoV-2 replication. Data was processed using Scanpy v1.5.1 [52], doublets were detected with scrublet v0.2.1 [53] and removed, ribosomal genes were removed and multisample integration was performed with BBKNN v1.3.12 [54]. Gene set enrichment analysis was performed with signatures retrieved from gsea-msigdb.org website [55] using following terms: HALLMARK_INTERFERON_GAMMA_RESPONSE M5913, HALLMARK_INTERFERON_ALPHA_RESPONSE M5911. Computations were automated with snakemake v5.5.4 [56].

**Deconvolution of bulk RNA-seq alveolar macrophage signatures** was performed using AutoGeneS v1.0.3[57] and signatures derived from integrated single cell RNA-Seq object. For details of the functions used and their parameters see the code https://github.com/NUPulmonary/2020_Grant.

### Statistical analysis

All statistical analysis was performed using R version 3.6.3. For all comparisons, normality was first assured using a Shapiro-Wilk test and manual examination of distributions. For parameters exhibiting a clear lack of normality, nonparametric tests were employed. In cases of multiple testing, P-values were corrected using false-discovery rate (FDR) correction. Adjusted P-values < 0.05 were considered significant.

### Visualization

All plotting was performed using ggplot2 version 3.3.1, with the exception of heatmaps, which were generated using pheatmap version 1.0.12. Sankey/Alluvial plots were generated using ggalluvial version 0.12.0[58]. Figure layouts were generated using patchwork version 1.01 and edited in Adobe Illustrator 2020.

## Data and code availability

Bulk RNAseq: Counts tables and metadata are included as Supplemental data files 2 and 3.

Single-cell RNA-seq: Counts tables and integrated objects are available through GEO with accession number GSE155249. Raw data is in the process of being deposited to SRA/dbGaP.

Code: https://github.com/NUPulmonary/2020_Grant.

## Tables and Supplementary Materials

### Supplementary Data Files

Supplementary Data File 1: Data related to k-means clustering and differential expression analysis (DEA) of bulk RNA-seq samples: gene cluster assignments, cluster GO enrichment, pairwise DEA across diagnosis. Excel workbook, multiple tabs.

Supplementary Data File 2: Raw bulk RNA-seq gene counts. Excel workbook. Supplementary Data File 3: Bulk RNA-seq metadata. Excel workbook.

Supplementary Data File 4: Marker genes for integrated scRNA-seq object containing cells from 5 patients with COVID-19. Excel workbook.

Supplementary Data File 5: Marker genes for integrated scRNA-seq object containing cells from 5 patients with COVID-19 and a single patient with bacterial pneumonia. Excel workbook.

Supplementary Data File 6: Differentially expressed genes between TRAM1 and TRAM2 clusters. Excel workbook.

Supplementary Data File 7: Standardized tool for pneumonia category adjudication.

Supplementary Data File 8: List of all SCRIPT study investigators, their affiliations and ORCID IDs.

---

**Supplemental Figure S1:**
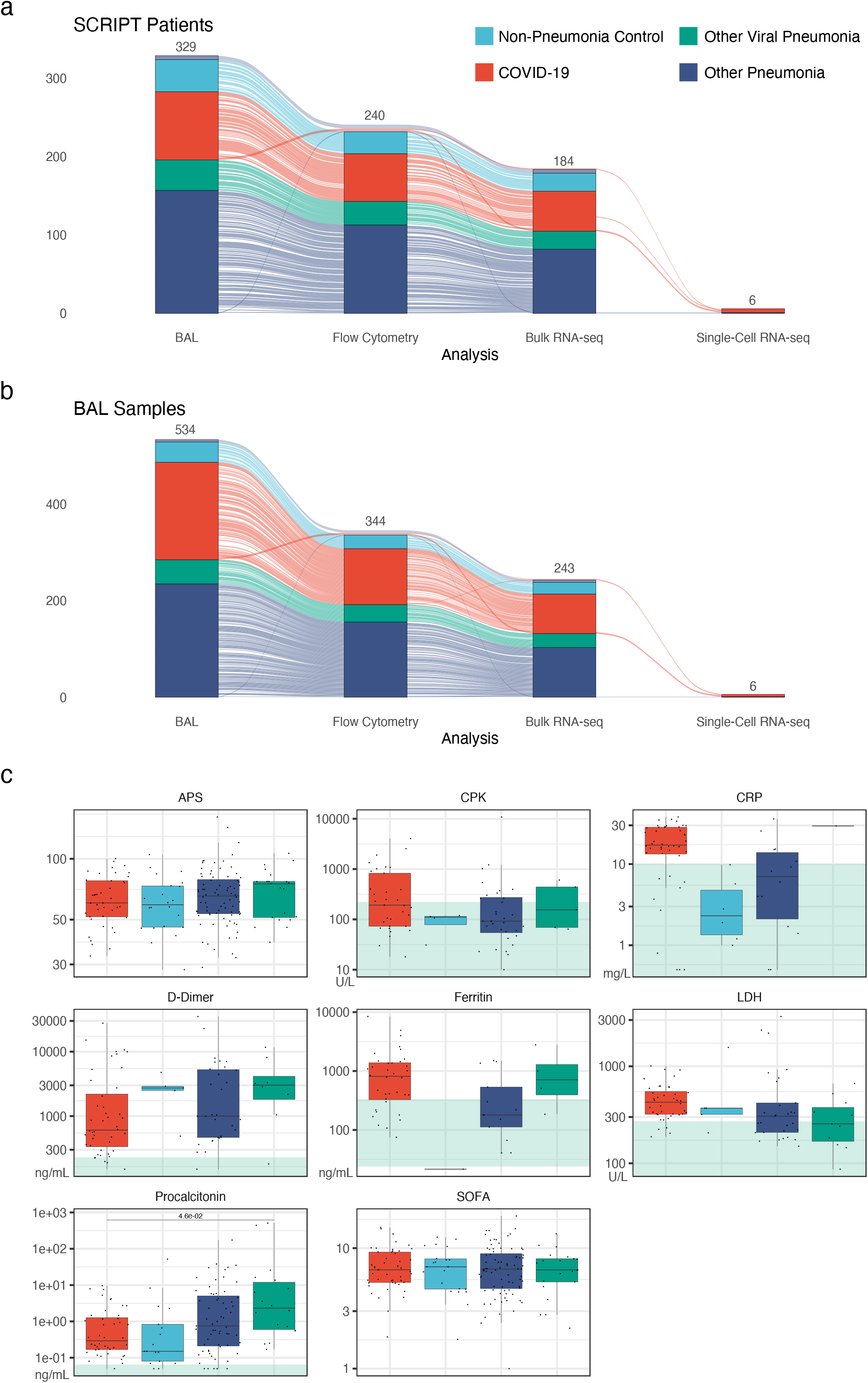
Overview of the study and biomarkers. **a**. Sankey diagram illustrating relationship between number of participants with COVID-19, other viral pneumonia, non-viral pneumonia (other pneumonia) and non-pneumonia controls 1) enrolled in the SCRIPT study (329 patients), 2) analyzed via flow cytometry (240 patients), 3) bulk RNA-seq on flow-sorted alveolar macrophages (184 patients) and 4) single-cell RNA-seq (6 patients). Some samples were cryopreserved and sorted post-cryorecovery. Since cryopreservation affects the number of granulocytes, these samples were not included in flow cytometric analysis, but were used for bulk RNA-seq profiling of flow-sorted alveolar macrophages. Skipped analyses are represented by alluvia flowing over the grouping bars. **b**. Sankey diagram illustrating relationship between number of BAL samples from participants with COVID-19, other viral pneumonia, non-viral pneumonia (other pneumonia) and non-pneumonia controls 1) enrolled in the SCRIPT study (534 samples), 2) analyzed via flow cytometry (344 samples), 3) bulk RNA-seq on flow-sorted alveolar macrophages (243 samples) and 4) single-cell RNA-seq (6 samples). **c**. The Sequential Organ Failure Assessment (SOFA) score, the Acute Physiology Score (APS) and inflammation biomarkers: creatine phosphokinase (CPK), C-reactive protein (CRP), D-dimer, ferritin, lactate dehydrogenase (LDH), procalcitonin (q = 4.6×10^−2^, pairwise Wilcoxon rank-sum tests with FDR correction). Green-shaded area indicates normal range.

**Supplemental Figure S2:**
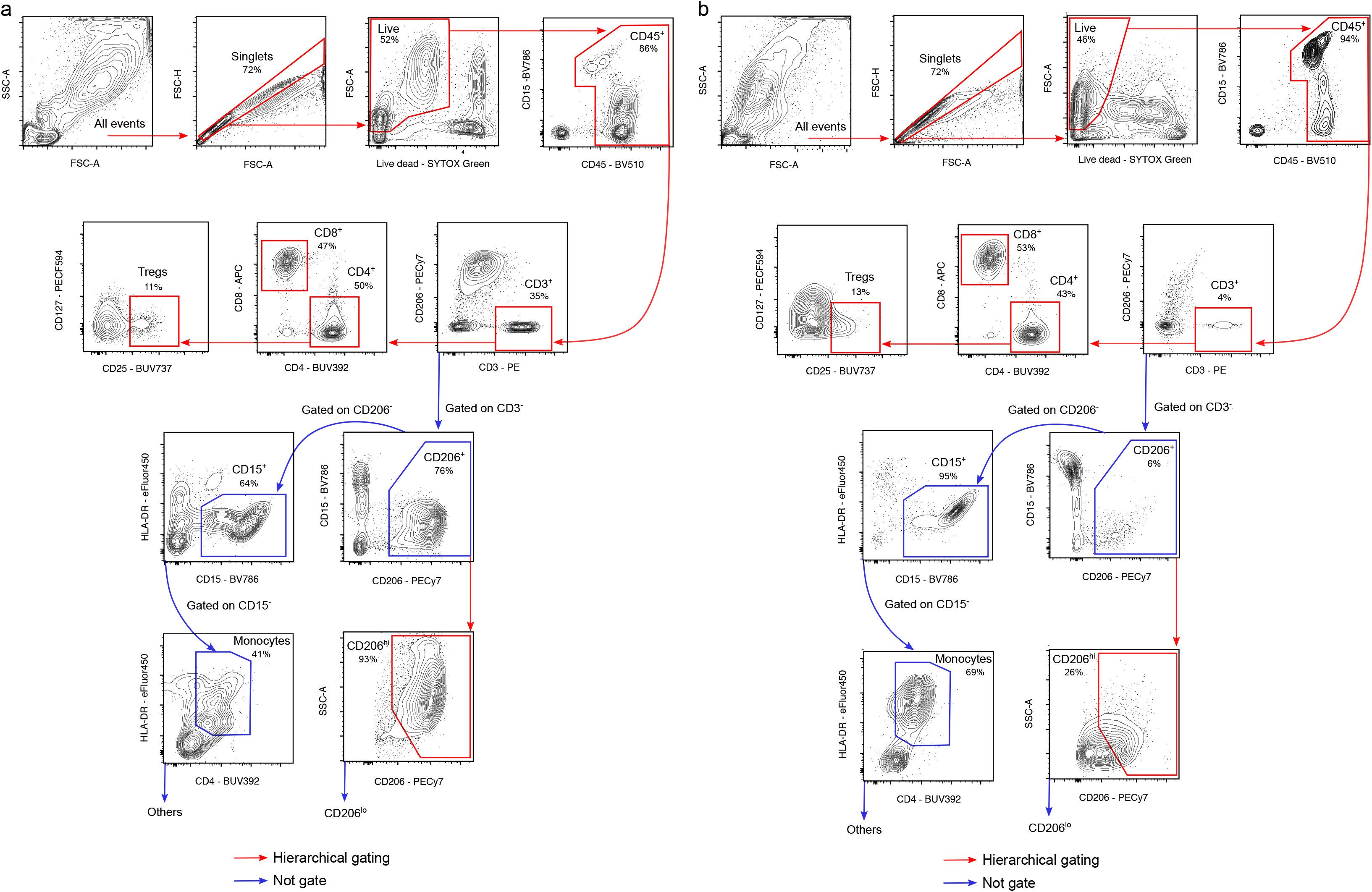
Representative gating strategy used to identify immune cell subsets in BAL samples. **a**. Representative sample from a patient without neutrophilia. **b**. Representative sample from a patient with neutrophilia and loss of CD206^hi^ alveolar macrophages. T cells were identified as CD3-positive and further subdivided into CD4+ and CD8+ T cells. Tregs were identified as CD3+CD4+CD25+CD127-. Neutrophils were identified as CD15+ cells. Alveolar macrophages were identified as CD206+ and further subdivided into CD206^hi^ and CD206^lo^ cells. Monocytes were identified as HLA-DR+CD4+ cells.

**Supplemental Figure 3.**
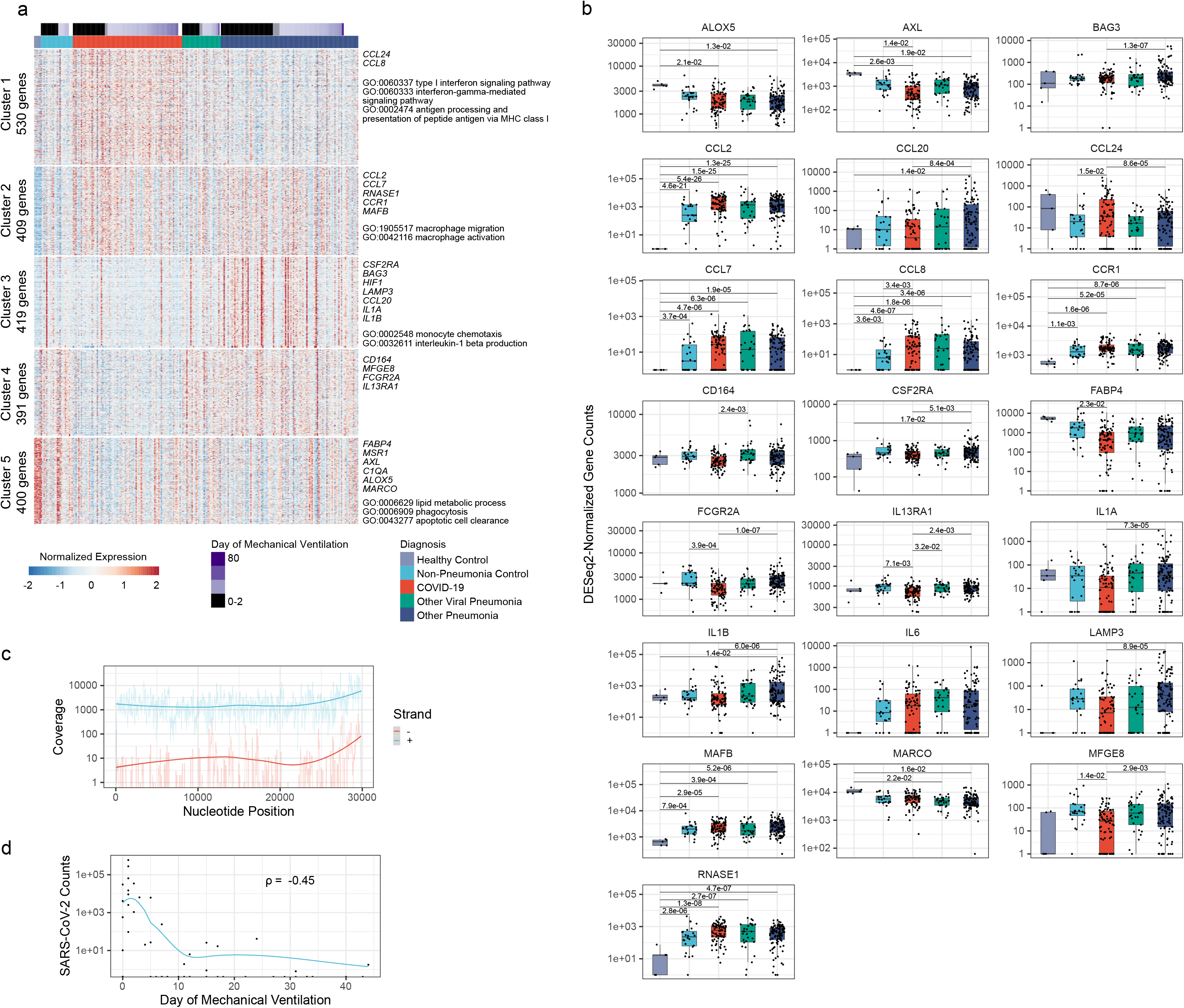
Transcriptomic analysis of alveolar macrophages identifies an interferon response signature as a feature distinguishing SARS-CoV-2 pneumonia from other types of pneumonia. **a**. k-means clustering of the 2,149 most variable genes across diagnoses using a likelihood-ratio test (LRT), columns represent each individual patient, grouped by diagnosis and ordered by intubation day. Representative genes and biological GO processes are shown for each cluster. Column headers are color-coded by the diagnosis and duration of mechanical ventilation. **b**. Expression of selected genes between the groups. Note that expression of *IL6* is not increased in any group. All significant comparisons are shown (q < 0.05, Wald test with FDR correction in DESeq2). **c.** Coverage plot of RNA-seq reads aligned to the SARS-CoV-2 genome. **d**. Correlation between detection of SARS-CoV-2 reads and disease progression (ρ = −0.45, P = 8.3×10^−4^, Spearman correlation).

**Supplemental Figure 4.**
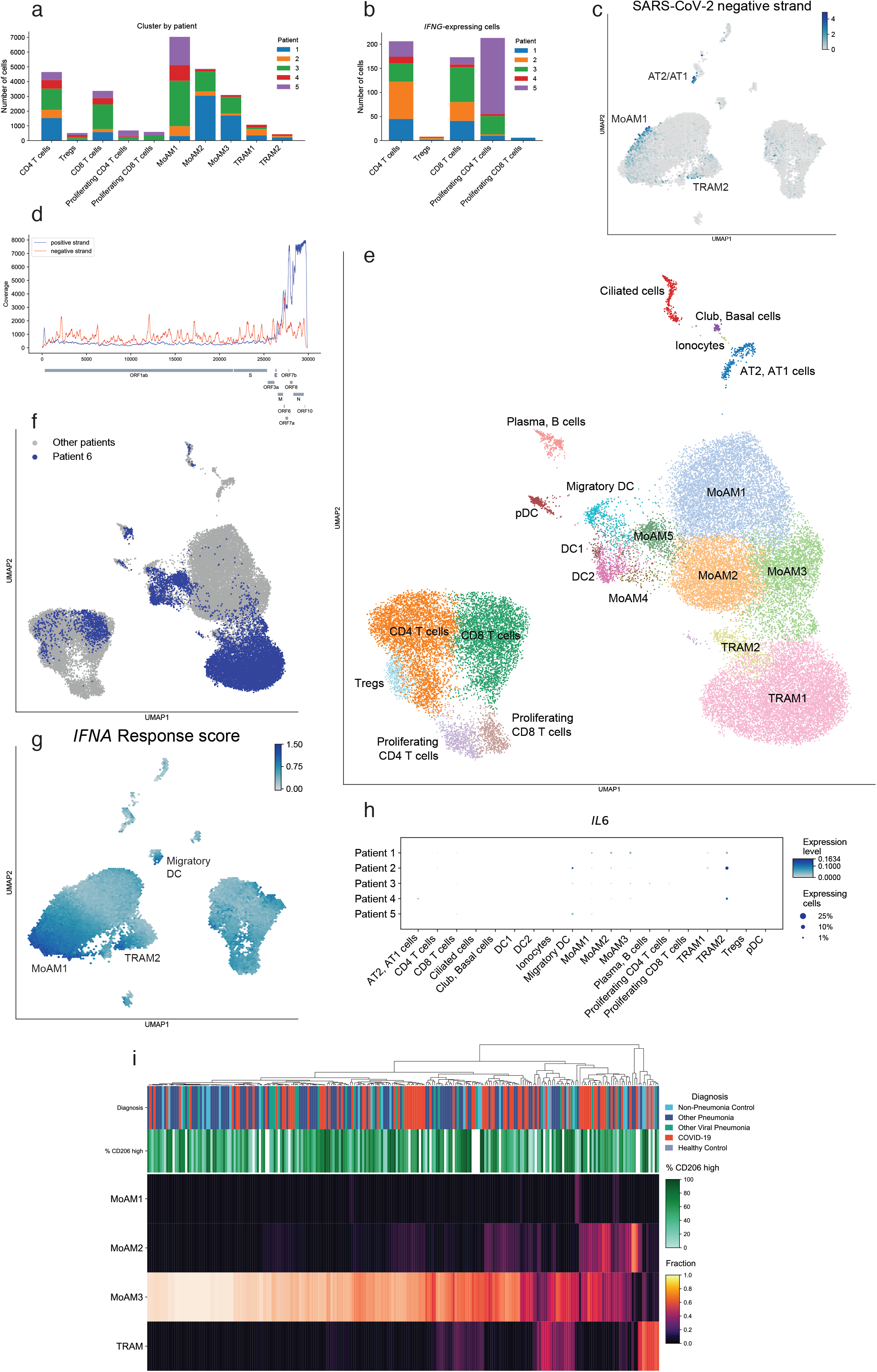
Single cell RNA-Seq identifies a positive feedback loop between IFNγ-producing T cells and SARS-CoV-2-infected alveolar macrophages. **a**. Subsets of alveolar macrophages and T cells are represented by the cells from all 5 patients with COVID-19. **b**. *IFNG* expression is detected in T cells from all 5 patients with COVID-19, T cells with at least one count of *IFNG* were used for analysis. **c**. Detection of the SARS-CoV-2 negative strand. Density projection plots, expression averaged within hexagonal areas on UMAP. **d**. Coverage plot of single cell RNA-seq reads aligned to SARS-CoV-2 genome. Cumulative data from 5 patients. Reads were aligned to genes on the positive strand or to the entire negative strand. **e**. UMAP plot showing integrative analysis of 39,331 cells isolated from 5 patients with severe COVID-19 within 48 hours after intubation (see **Figure 4**) and a single intubated patient with bacterial pneumonia. AT1 – alveolar epithelial type 1 cells; AT2 – alveolar epithelial type 2 cells; DC1 – conventional dendritic cells type 1, *CLEC9A*+; DC2 – conventional dendritic cells type 2, *CD1C*+; Migratory DC – migratory dendritic cells, *CCR7*+; pDC – plasmacytoid dendritic cells, *CLEC4C*+; TRAM – tissue-resident alveolar macrophages; MoAM – monocyte-derived alveolar macrophages; Tregs – regulatory T cells, *FOXP3*+. **f.** UMAP plot showing cells from a patient with bacterial pneumonia (Patient 6) from the integrative analysis on Figure S4e. **g**. Density projection of hallmark interferon-alpha response genes among 5 patients with COVID-19 on top of UMAP. **h**. Dot plot showing *IL6* expression across the cell types split by patient. Dot size is proportional to the number of cells expressing *IL6* in the corresponding cluster. **i**. Heatmap demonstrating proportion of specific cell types predicted from deconvolution of the transcriptomic profiles from flow-sorted alveolar macrophages based on signatures from integrated single-cell RNA-seq analysis.

