## Supplementary Data File 7 for "Alveolitis in severe SARS-CoV-2 pneumonia is driven by self-sustaining circuits between infected alveolar macrophages and T cells"

### Pneumonia Episode Category Assessment

Record ID

______________________

Is this assessment for the initial pneumonia episode?

Instructions: If there is more than one episode, select "No" and other episodes will be created according to research coordinator definitions. Please review to make sure that subsequent episode is distinct from the initial. If not, please ask Helen or Nicole to delete &/or revise episode to correspond to correct BAL time point.

◯ Yes
◯ No

If not, is this the final pneumonia episode?

Instructions: Select "Yes" if this is the last episode for current hospitalization. Select "No" if more than two episodes.

◯ Yes
◯ No

Assessment date:

______________________
(Note: Day of Initial BAL or new HAP/VAP episode.)

Study day (auto-calculated):

______________________

This patient is immunocompromised.

Note: Data from Demographics form. If this is incorrect, please change in Demographics form.

This patient is NON immunocompromised.

Note: Data from Demographics form. If this is incorrect, please change in Demographics form.

If no, study day (this assessment is for succeeding pneumonia episodes): ______________________

Category of patient:

Instructions: Using standard 48 hour rules for CAP vs HAP (hospital admission), HAP vs VAP (ventilation).

Two critical issues:
Default is pneumonia - have subsequent indeterminate option if very equivocal.
No pneumonia - review carefully to assess whether there is good evidence that pneumonia is not present. Will need to fill in subsequent questions about evidence.

◯ Clinical CAP (not hospitalized within the last 7 days)
◯ Clinical HAP (current admission >48 hours or discharged from a healthcare facility within the last 7 days where admission >24 hours)
◯ Clinical VAP (on ventilator >48 hours or reintubated < 24 hours from extubation)
◯ Non-pneumonia control

| | |
|---|---|
| If Clinical CAP: | ◯ Etiology defined<br>◯ Viral NP/OP only<br>◯ Culture-negative |
| If culture-negative: | ◯ Immunocompromised<br>◯ Nonimmunocompromised<br>(Note: Please select an option. This does not auto-populate.) |
| If Clinical HAP: | ◯ Etiology defined<br>◯ Culture-negative (%PMNs > 50%)<br>◯ Culture-negative (%PMNs < 50%)<br>◯ Viral only<br>◯ Indeterminate |
| If culture-negative (%PMNs > 50%): | ◯ Immunocompromised<br>◯ Nonimmunocompromised |
| If culture-negative (%PMNs < 50%): | ◯ Neutropenic (ANC < 500/uL)<br>◯ Immunocompromised<br>◯ Nonimmunocompromised |
| If Clinical VAP: | ◯ Etiology defined<br>◯ Culture-negative (%PMNs > 50%)<br>◯ Culture-negative (%PMNs < 50%)<br>◯ Viral only<br>◯ Indeterminate |
| If culture-negative (%PMNs > 50%): | ◯ Immunocompromised<br>◯ Nonimmunocompromised |
| If culture-negative (%PMNs < 50%): | ◯ Neutropenic (ANC < 500/u)<br>◯ Immunocompromised<br>◯ Nonimmunocompromised |
| If non-pneumonia control, cause of infiltrate: | ◯ ARDS<br>◯ Aspiration<br>◯ Atelectasis<br>◯ Fibrosis<br>◯ Fluid overload<br>◯ Heart failure/pulmonary edema<br>◯ Pleural effusion<br>◯ Pulmonary hemorrhage<br>◯ Other (please specify)<br>◯ Unknown |
| Please specify: | ______________________ |
| If non-pneumonia control: | ◯ Infection<br>◯ Known condition<br>◯ Unknown |

| | |
|---|---|
| If infection: | ◯ Cholangitis/cholecystitis<br>◯ Colitis<br>◯ Other intra-abdominal<br>◯ Line infection<br>◯ Tracheobronchitis<br>◯ Urinary tract<br>◯ Wound/skin<br>◯ Other (please specify) |
| Please specify: | ______________________ |
| If known condition, cause of fever/leukocytosis: | ◯ Aspiration<br>◯ Atelectasis<br>◯ Drug fever<br>◯ Pancreatitis<br>◯ Other (please specify)<br>◯ None<br>◯ Unknown |
| Please specify: | ______________________ |

Change form status to "complete" when finalized.
